# Heterotrimeric G protein αi2 Sequesters RasGAP to Control Neutrophil Sensitivity and Chemotaxis

**DOI:** 10.64898/2026.05.01.722239

**Authors:** Xuehua Xu, Woo Sung Kim, Arthur Lee, Riley Kim, Caleb Zhao, Huie Jing, Helen Su, Tian Jin

## Abstract

G protein–coupled receptors (GPCRs) direct neutrophil chemotaxis through heterotrimeric G proteins, yet the downstream effectors of the predominant G*a*i isoform, G*a*i2, remain incompletely defined. Here we identify the Ras GTPase-activating protein CAPRI (RASA4) as a functional effector of G*a*i2 that links GPCR signaling to Ras adaptation. Using AlphaFold3-based structural modeling and binding free-energy calculations and experimental verification, we reveal that constitutively active and structurally altered G*a*i2 mutants (Q205L and T182A) exhibit enhanced binding to CAPRI. Neutrophils expressing these mutants display elevated basal Ras activity, heightened sensitivity to chemoattractant, and improved chemotaxis in low- or subsensitive-concentration gradients. However, these cells exhibit excessive Ras activation and impaired chemotaxis at high, saturating chemoattractant concentrations, while maintaining near-normal responses at intermediate concentrations. These results reveal an upward shift in the concentration range for efficient chemotaxis. Our findings not only establish a previously unrecognized G*a*i2–CAPRI signaling axis that tunes Ras adaptation but also define a mechanism by which heterotrimeric G proteins calibrate leukocyte navigation across diverse chemoattractant gradients.

## INTRODUCTION

Cells sense diverse extracellular stimuli—including hormones, ions, small molecules, and light—through G protein–coupled receptors (GPCRs), which transmit signals to intracellular effectors to generate appropriate cellular responses (Neves et al., 2002). Upon ligand binding, GPCRs activate associated heterotrimeric G proteins by promoting GDP–GTP exchange on the Ga subunit. Activated Ga dissociates from GBy to regulate downstream effectors, and intrinsic GTPase activity of Ga subsequently hydrolyzes GTP to GDP, thereby terminating signaling and permitting reassociation of the heterotrimer. The timing and magnitude of G protein signaling are thought to be tightly controlled by regulators of G protein signaling (RGS) proteins, which accelerate GTP hydrolysis and promote signal deactivation (Masuho et al., 2020). Several direct effectors of GBy, including phospholipase C B2/3 (PLCB2/3), phosphoinositide 3-kinase y (PI3Ky), P-Rex1, and Elmo, mediate key signaling pathways controlling neutrophil chemotaxis (Kuang et al., 1996; Welch et al., 2002; Suire et al., 2006). Adenylyl cyclase (ACA) is the known direct effector of Gαi1 subunits (Dessauer et al., 1998; Taussig et al., 1993). Gai2 is the predominant Gai subunit in neutrophils and is essential for chemoattractant signaling (Kuwano et al., 2016; Yan et al., 2021). However, the downstream effectors of Gαi2 remain poorly defined.

Ras plays a central role in chemotaxis of eukaryotic cells. Ras is a molecular switch that is activated by guanine exchange factors (GEFs) and deactivated by GTPase-activating proteins (GAPs). CAPRI (Calcium-promoted Ras inactivator, also known as RASA4) is a multi-domain signaling protein containing tandem C2 domains, a pleckstrin homology (PH) domain, and a Ras GTPase-activating protein (GAP) domain. CAPRI integrates calcium and membrane signals to regulate small GTPase activity and signal transduction spatiotemporally (Lockyer et al., 2001). We recently found that CAPRI is highly expressed in human neutrophils, mediates Ras adaptation, and plays an essential role in chemotaxis (Xu et al., 2021b). Emerging evidence suggests that RasGAPs interact functionally with heterotrimeric G proteins, including Gai2 in human T cells and Gα2 in the model organism *Dictyostelium discoideum* (Ham et al., 2024b; Xu et al., 2026). However, the structural basis and nucleotide dependence of heterotrimeric G protein−RasGAP interaction remain poorly defined. In this study, we used AlphaFold 3–based structural modeling and binding free-energy calculations to characterize the molecular architecture of CAPRI in complex with Gai2 (WT, Q205L, and T182A) in both GDP- and GTP-bound states. By integrating structural modeling with experimental observations, we define the nucleotide- and mutation-dependent determinants governing CAPRI–Gai2 interactions. Furthermore, we characterize the effect of these interactions on sensitivity and chemotaxis in human neutrophils. We found that neutrophils expressing these mutants display elevated basal Ras activity, heightened sensitivity to chemoattractant, and improved chemotaxis in low- or subsensitive-concentration gradients. However, these cells exhibit excessive Ras activation and impaired chemotaxis at high, saturating chemoattractant concentrations, while maintaining near-normal responses at intermediate concentrations. These findings revise the current understanding of how G protein and Ras signaling intersect to fine-tune chemotactic responses, providing a new framework for investigating leukocyte navigation in complex tissue environments.

## RESULTS

### Constitutively Active G*a*i2 Mutants Exhibit Enhanced Binding to CAPRI Upon Chemoattractant Stimulation

Primary human (Hs) and mouse (Mm) neutrophils express multiple Gα subunits that play essential roles in several functional processes (**Figure 1A**). Human neutrophil-like (HL60) cells, a widely used model for neutrophil studies (Birnie, 1988; Houk et al., 2012; Rincon et al., 2018), exhibit a similar Ga subunit expression pattern upon differentiation (**Figure 1B**), consistent with a previous report (Rincon et al., 2018). We previously identified CAPRI as a key Ras GTPase-activating protein (GAP) that mediates deactivation of Ras for proper adaptation and chemotaxis in HL60 cells (Xu et al., 2021b). Notably, CAPRI interacts with multiple α subunits of G protein, including Gαi2− a critical mediator of neutrophil chemotaxis (**Figure 1C**). Recent findings have shown that constitutive-active (CA) mutants of Gai2 can sequester RASA2, another Ras GAP, leading to Ras hyperactivation in T cells (Ham et al., 2024b). We found that Gαi2 and Ras share the same interaction domain on CAPRI, suggesting a potential competitive binding (**Figure 1C**). To investigate this further, we examined Ras GAP proteins that interacts with Gαi2 in HL60 cells expressing GFP-tagged WT-Gαi2, Gαi2-Q205L, or Gαi2-T182A (**Figure S1**) following chemoattractant N-Formylmethionyl-leucyl phenylalanine (fMLP) stimulation, using co-immunoprecipitation (co-IP). In agreement with previous findings (Ham et al., 2024a), we detected a weak interaction between RASA2 and two active mutants of Gαi2. Both Gαi2-Q205L and Gαi2-T182A showed a predominant interaction with CAPRI upon stimulation (**Figure 1D**). Consistently, a higher level of CAPRI expression than that of RASA2 provides an explanation for this stronger binding between CAPRI and Gαi2, notably in the case of overexpression (**Figure S2**). Taken together, our results indicate that Gαi subunits directly associate with CAPRI and that constitutive mutants of Gαi2 display a stronger binding with CAPRI.

**Figure 1.**
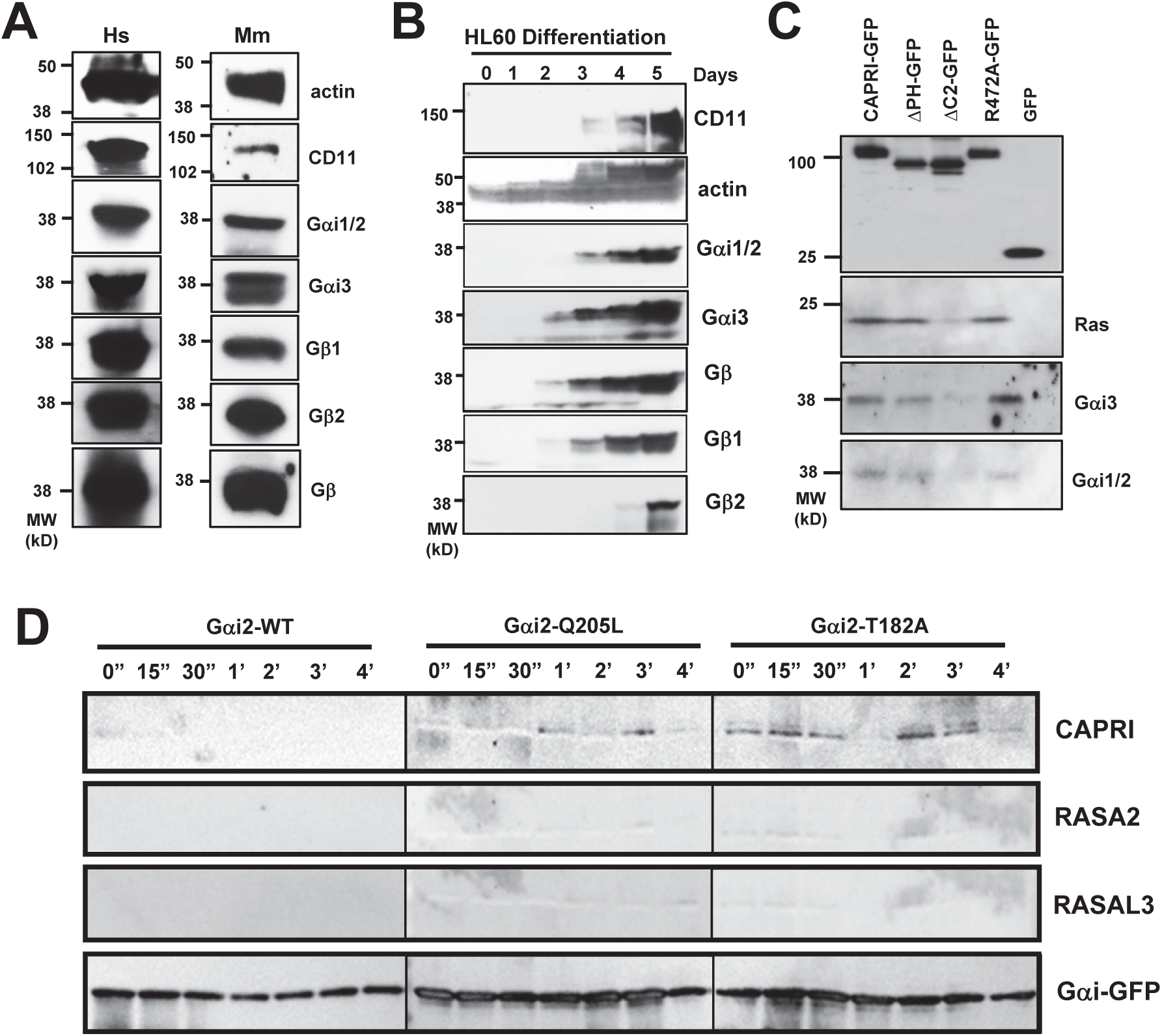
Constitutive-active (CA) mutants of Gαi2 display a higher binding affinity toward CAPRI. **A.** Multiple heterotrimeric G proteins are expressed in primary human (Hs) or mouse (Mm) neutrophils. **B.** The expression pattern of heterotrimeric G proteins during the differentiation of human neutrophil-like HL60 cells. **C.** CAPRI interacts with multiple Gαi subunits determined by co-immunoprecipitation (co-IP) assay. HL60 cells expressing wild-type (WT) or mutants of CAPRI tagged with turbo-GFP were stimulated with saturating (10 μM) fMLP at time 0 s. **D.** Dynamic interaction between Gαi2 (WT and constitutive-active mutants) and primary Ras GAPs in HL60 cells upon fMLP stimulation, as determined by co-immunoprecipitation (co-IP). Aliquots of cells expressing wild-type (WT) or CA mutants of Gαi2 at the indicated time points were taken and subjected to the co-IP assay and the indicated proteins were detected by western using their specific antibodies.

### Validation of the structures of G*a*i2 WT and Mutants simulated using AlphaFold3

To gain mechanistic insight into constitutive active mutants of Gαi2, we modeled the structures of Gαi2 WT and Gαi2 mutants Q205L and T182A using AlphaFold3 (**Figure 2**). We first predicted the structures of Gai2-WT in its GDP- and GTP-bound forms with high confidence. Gai2-WT adopts the canonical architecture composed of a Ras-like GTPase domain and an a-helical domain, separated by a deep cleft that accommodates GDP or GTP (**Figure S3A**). Consistent with previous structural studies of the related Gαi1 (Wall et al., 1995), the aN helix at the N-terminus is positioned to interact with the GBy subunits, whereas the C-terminal a5 helix forms the primary interface with activated GPCRs (Rasmussen et al., 2011). Comparison of GDP- and GTP-bound WT models revealed substantial conformational rearrangements involving the N-terminus, Switch I–III regions, and the C-terminal a5 helix (**Figure 2A**; **Video S1**), consistent with the previous report (Lambright et al., 1994). Predicted complexes of Gai2 with GB1, GB2, and GBy preserved the conserved heterotrimeric architecture as reported previously (**Figure S3B**) (Lambright et al., 1996; Wall et al., 1995).

**Figure 2.**
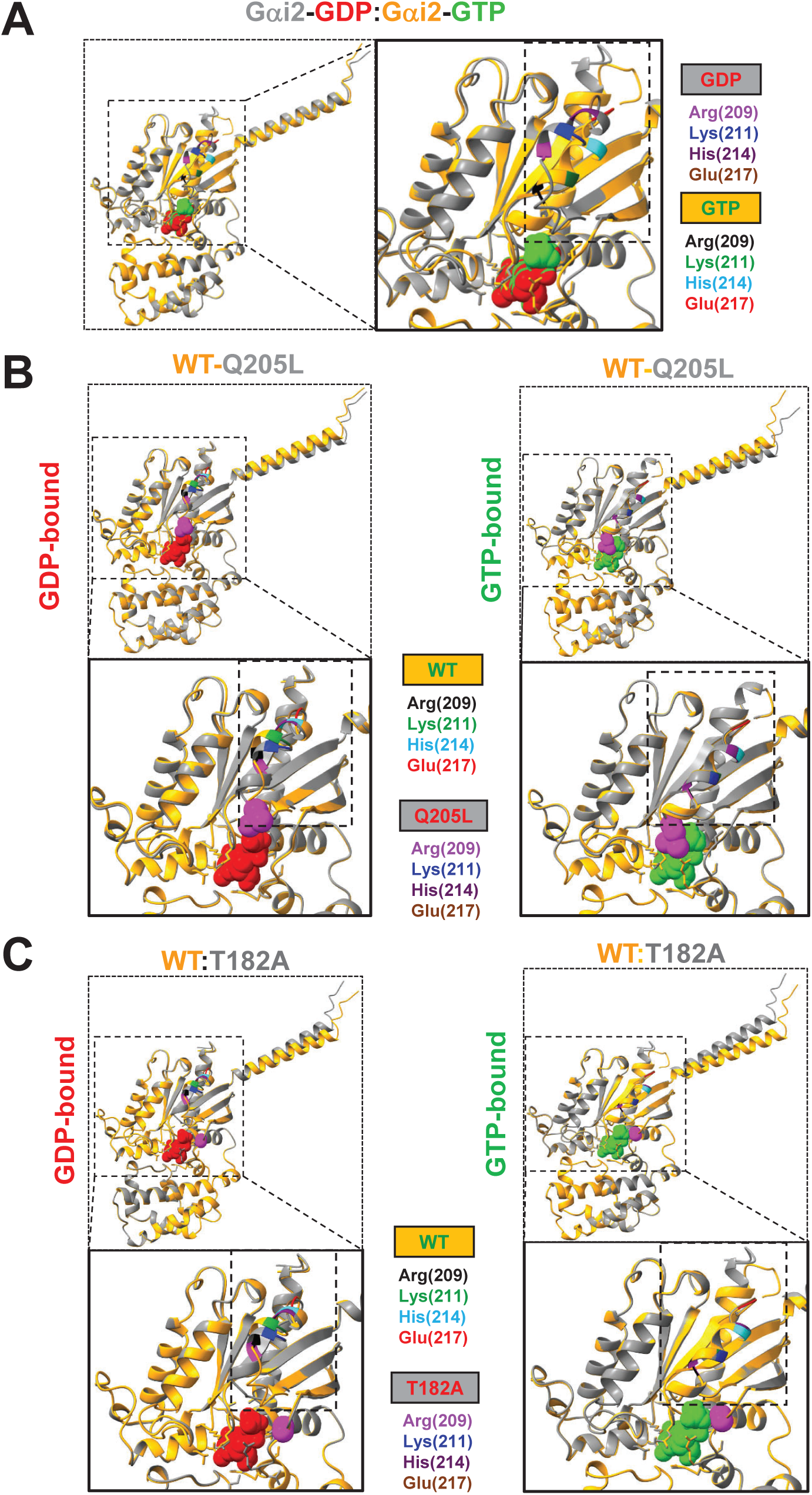
Validation of the structures of G*a*i2 WT and Mutants simulated using AlphaFold3. **A.** Alignment of Gai2–GDP (gray) and Gai2–GTP (gold) structures. GDP is shown as red spheres, whereas GTP is shown as green spheres. The switch region is enlarged in the right panel to illustrate conformational differences between the nucleotide-bound states. The enlarged view emphasizes structural rearrangements within the switch region. Key residues exhibiting significant positional shifts between the GDP- and GTP-bound forms are shown in the corresponding colors. A complete set of structures is provided in **Video S1**. **B.** Structural alignment of Gai2–WT and Gai2–Q205L. Superposition of Gai2–WT (gold) and Gai2–Q205L (grey) in complex with GDP either GDP (left; GDP, red spheres) or - GTP (right; GTP, green spheres). The switch region is enlarged in the lower panel to highlight conformational differences between WT and mutant structures. The Q205L substitution is shown in magenta. Key residues in Switch II are labeled with residue numbers and are highlighted in the same color scheme in the structure. A complete set of structures is provided in **Video S2**, top panel (GDP-bound, left; GTP-bound, right). **C.** Structural alignment of Gai2–WT and Gai2–T182A. Superposition of Gai2–WT (gold) and Gai2–T182A (grey) in complex with GDP either GDP (left; GDP, red spheres) or - GTP (right; GTP, green spheres). The switch region is enlarged in the lower panel to highlight conformational differences between WT and mutant structures. The T182A substitution is shown in magenta. Key residues in Switch II are labeled with residue numbers and are highlighted in the same color scheme in the structure. A complete set of structures is provided in **Video S2**, bottom panel (GDP-bound, left; GTP-bound, right).

We next modeled the Q205L and T182A mutants. Q205, located within the catalytic Switch II region, is essential for GTP hydrolysis, whereas T182 lies adjacent to the nucleotide-binding pocket within the catalytic core (**Figure S3A**, middle and right panels) (Coleman et al., 1994; Hewitt et al., 2023). In the GDP-bound state, the Q205L mutation induced pronounced conformational rearrangements in Switch II, particularly around Arg209, Lys211, His214, and Glu217, accompanied by changes in the C-terminus. By contrast, the GTP-bound Q205L structure showed minimal alterations in Switch II but retained differences at the C-terminus (**Figure 2B**; **Video S2**, top panel). The T182A mutant displayed distinct structural effects. In the GDP-bound state, T182A produced conformational shifts in Switch II together with more pronounced changes at the C-terminus. In the GTP-bound state, T182A induced substantial rearrangements in both the N- and C-terminal regions, while Switch II remained largely unchanged (**Figure 2B**; **Video S2**, bottom panel). Together, these structural analyses suggest that Q205L and T182A promote constitutive activity through distinct mechanisms, with Q205L primarily perturbing the catalytic Switch II network and T182A altering terminal domain dynamics that may influence receptor and heterotrimer interactions.

### A stronger binding between G*a*i2 mutants and CAPRI simulated by AlphaFold3

To gain mechanistic insight into the dynamic interaction between CAPRI and Gai2 (WT or mutants), we next modeled structures of CAPRI and their binding properties using AlphaFold3. The predicted CAPRI structure revealed the expected domain organization, including the C2A, C2B, GAP, and PH domains, each modeled with high confidence (**Figure 3B**, left panel). Notably, the relative orientation of the C2B and GAP domains closely matches a recently reported structure (**Figure 3A**; **Figure 3B**, right panel; Paul et al., 2025), supporting the reliability of the predicted model. We next modeled complexes between CAPRI and Gai2 in both GDP- and GTP-bound states (**Figure 3C**; **Video S3**). In both nucleotide-bound states, the C2 and GAP domains of CAPRI engage the Ras-like GTPase domain of Gai2-WT (**Figure S4A**). Binding free-energy calculations yielded predicted values of -13.2 kcal/mol for CAPRI–Gai2–GDP and -12.7 kcal/mol for CAPRI–Gai2–GTP, corresponding to inferred dissociation constants of approximately 2 × 10-^10^ M and 5.0 × 10-^10^ M, respectively (**Figure S4B**). Because these values are derived from computational simulations, they should be interpreted qualitatively rather than quantitatively. Nevertheless, the results suggest moderately weaker binding in the GTP-bound state, consistent with the trend observed experimentally (**Figure 1C**).

**Figure 3.**
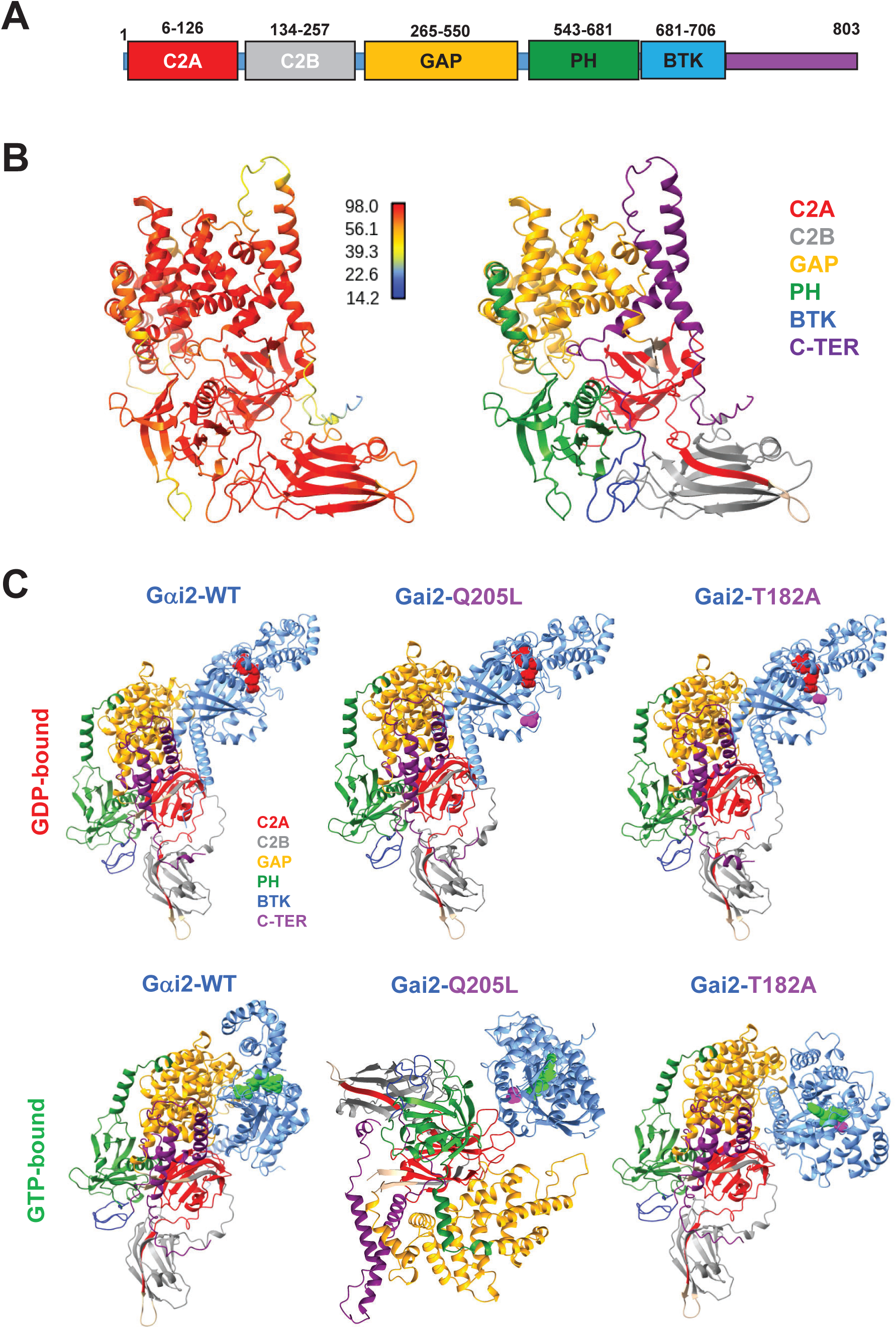
A stronger binding between G*a*i2 mutants and CAPRI simulated by AlphaFold3. **A.** A schematic diagram illustrating the domain composition of human CAPRI, with domain names and corresponding residue ranges indicated. **B.** Predicted structure of CAPRI. The AlphaFold 3–predicted structure is shown on the left, colored by confidence score (pLDDT), and the domain architecture of CAPRI is shown on the right. CAPRI domains are color-coded as follows: C2A (red), C2B (grey), RasGAP (gold), PH (green), BTK (dark blue), and C-terminus (dark magenta). **C.** Predicted CAPRI–Gai2 interactions. Structural models depict CAPRI in complex with WT-, Q205L-, or T182A-Gai2 in the GDP (red spheres)- or GTP (green spheres)-bound state. CAPRI domains are color-coded as shown in **B**. Point mutations of Q205L and T182A are highlighted in magenta. See a complete set of the structures in **Video S3.**

We further modeled CAPRI interactions with the Gai2 mutants Q205L and T182A. The GDP-bound Gai2-Q205L and Gai2-T182A mutants displayed binding modes similar to WT. In the GTP-bound state, both mutants exhibited altered binding properties relative to WT (**Figure S4A**), suggesting that mutation-induced conformational changes influence CAPRI engagement. Notably, Gai2-T182A showed stronger predicted binding to CAPRI (**Figure S4B**), consistent with the experimental data (**Figure 1D**). Although the overall binding affinity of Gai2-Q205L was predicted to be slightly lower than that of WT, the Q205L–GTP complex displayed stronger binding than Q205L-GDP, again consistent with experimental observations. Together, these structural analyses suggest that CAPRI can engage both inactive and active conformations of Gai2, with constitutively active mutations altering the conformational landscape of the GTPase domain to modulate CAPRI binding dynamics and potentially stabilize signaling-competent complexes.

### Increased Ras Activation in Neutrophils Expressing Constitutively Active Gαi2 Mutants Upon Chemoattractant Stimulation

To assess the functional consequence of enhanced interaction between the Gαi2 mutants with CAPRI, we evaluated Ras activation using a pull-down assay in HL60 cells expressing Gαi2-WT, -Q205L, or -T182A following stimulation with 10 µM fMLP, as described previously (Xu et al., 2021b). Equal numbers of cells were collected and lysed on ice. In WT-Gai2 cells, fMLP stimulation induced rapid but transient Ras activation, followed by adaptation. By contrast, cells expressing Q205L or T182A displayed elevated basal Ras activity and sustained Ras activation after stimulation (**Figure 4A**). Densitometric analysis from three independent experiments (**Figure 4A** and **Figure S5**) confirmed these results. Ras activity was quantified by normalizing the signal intensity at time 0 s to 1, with subsequent values expressed as Ii/Io (**Figure 4B**).

**Figure 4.**
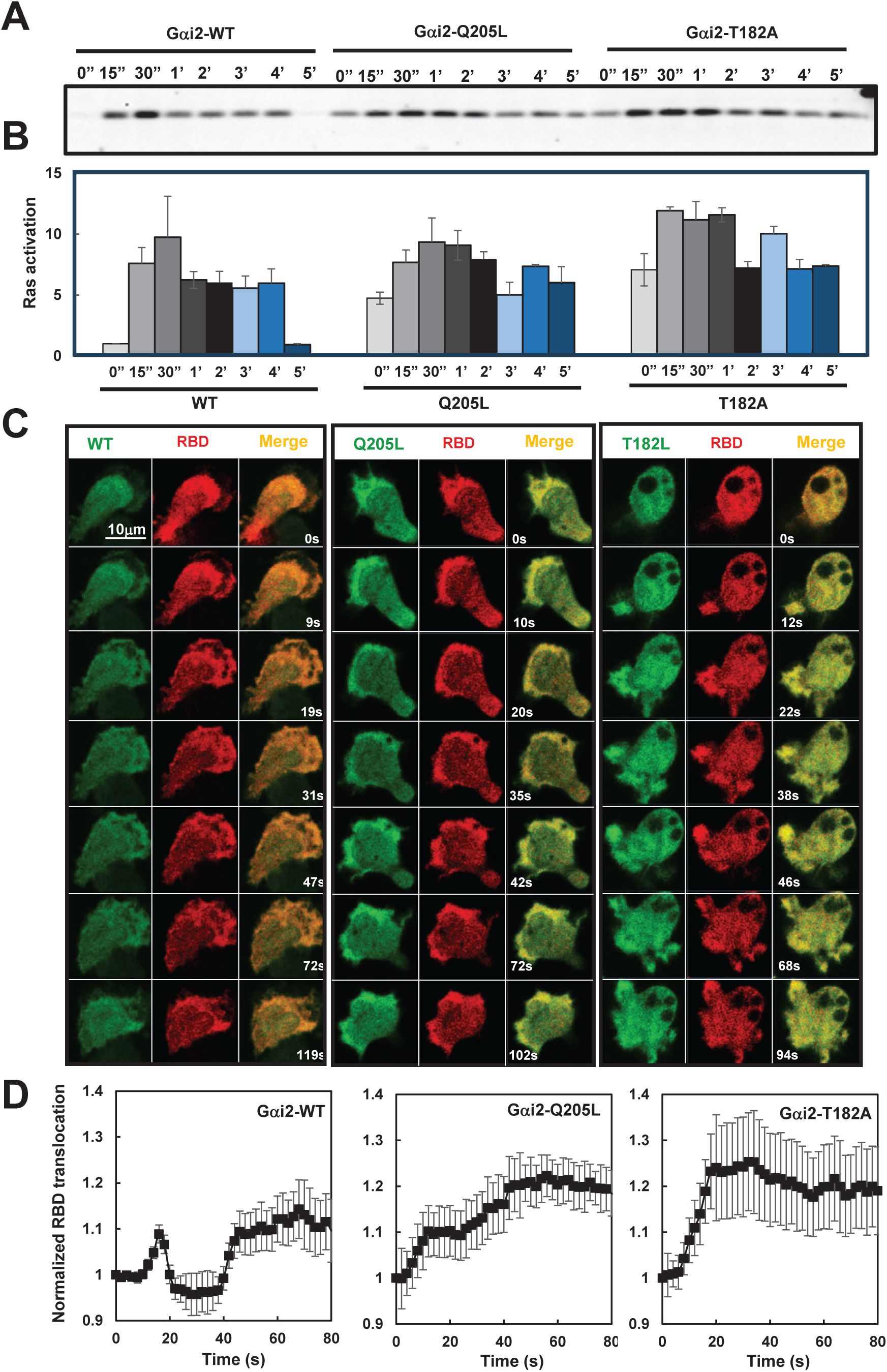
Increased Ras activation in neutrophils expressing CA-Gαi2 in response to chemoattractant stimulation. **A.** Ras activation in neutrophils expressing WT, Q205L, or T182A of Gai2 in response to saturating (10 mM) fMLP stimulation, as determined by a pull-down assay. Aliquots of cells were taken before or after fMLP stimulation at the indicated time points and subjected to the pull-down assay and subsequent western blot detection of active Ras as previously described. **B.** Normalized quantitative densitometry of the active Ras from three independent experiments, including the result presented in **A** and two sets of independent measurements shown in **Figure S5**. The intensity ratio of active Ras in WT at time 0 s was normalized to 1. Mean ± SD from the three independent experiments is shown. **C.** Visualization of Ras activation by plasma membrane (PM) translocation of active Ras biosensor, RBD-RFP, in the cells expressing either WT or CA mutants of Gai2 in response to saturating (10 mM) fMLP stimulation using confocal microscopy. Cells expressing either GFP tagged WT or CA-mutants of Gai2 (green) and RBD-RFP (red) were stimulated with fMLP right after the first frame. Arrows indicate the localization of RBD-RFP before or after stimulation. Scale bar = 10 mm. See **Video S4** (WT, top panels; Q205L, middle panels, and T182A, bottom panels) for complete set of cell responses. **D.** Quantitative measurement of Ras activation by PM translocation of RBD-RFP in the cells expressing either WT or CA-mutants of Gai2. To assess RBD-RFP plasma membrane translocation quantitatively, cytosolic depletion of CAPRI-GFP was measured. Regions of interest (ROIs) were selected in the cytoplasm and were tracked throughout the imaging period. Due to the migratory behavior of the cells, 5 cells of each group that exhibited typical responses with minimal movement were selected for quantitative analysis. Mean ± SD is shown.

To visualize Ras activation dynamics, confocal microscopy was performed using the active Ras biosensor RBD-RFP (red) in cells expressing WT, Q205L, or T182A Gai2 (green). WT-Gai2 cells exhibited transient plasma membrane recruitment of RBD-RFP, whereas Q205L- and T182A-expressing cells showed stronger and more sustained membrane localization, accompanied by expansion of the leading edge during cell migration (**Figure 4C**; **Video S3**). Quantification of RBD-RFP membrane translocation was performed by tracking cytoplasmic fluorescence depletion within selected regions of interest while avoiding the nucleus. Due to cell migration during imaging, 4–5 representative cells displaying typical responses with minimal movement were analyzed. Measurements across multiple cells confirm enhanced and prolonged Ras activation in Q205L- and T182A-expressing cells (**Figure 4D**). To assess membrane translocation quantitatively, we measured cytosolic depletion of CAPRI-GFP as previously reported (Xu et al., 2025; Xu et al., 2021b). Regions of interest (ROIs) were selected in the cytoplasm, and were tracked throughout the imaging period. The results shown in **Figure 4D** across multiple cells further confirmed the hyperactivation of Ras in Q205L or T182A Gαi2 cells. These findings suggest that constitutively active Gai2 mutants alter CAPRI-mediated regulation of Ras signaling, leading to impaired adaptation and sustained Ras activation following chemoattractant stimulation.

### Impaired Chemotaxis of Gαi2-Active Mutant-Expressing Neutrophils in Saturating Chemoattractant Gradients

Dynamic Ras activation and adaptation are critical for maintaining directional migration and persistent Ras signaling results in impaired neutrophil chemotaxis (Xu et al., 2021b). The prolonged Ras activation in Q205L- and T182A-expressing cells indicates impaired negative regulation of chemoattractant signaling. As expected, defective chemotaxis of Gαi2-active mutant-expressing neutrophils as has been reported previously (Ham et al., 2024b). To examine further the effects of Ras hyperactivation on neutrophil chemotaxis, we reanalyzed the migratory behavior of HL60 cells expressing endogenous Gai2 or overexpressing WT-, Q205L-, or T182A-Gai2 using DIAS software (Wessels et al., 1998). In gradients generated from a 1 µM fMLP source, cells expressing endogenous Gai2 or overexpressing WT-Gai2 exhibited comparable chemotactic performance, including directionality, migration speed, and total path length, although cells expressing endogenous Gai2 displayed slightly improved polarization (**Figure 5A**, left; **Video S4**; **Figure 5B**). By contrast, cells expressing the Q205L or T182A mutants showed a significant reduction in all measured chemotaxis parameters. Similar defects were observed in saturating gradients (1 µM) for LTB4 and SDF1a (**Figure 5A**, right; **Figure S6**), indicating that the impaired migration phenotype was not specific to a single chemoattractant. These results quantitatively demonstrate that constitutively active Gai2 mutants disrupt efficient neutrophil chemotaxis under saturating chemoattractant gradients.

**Figure 5.**
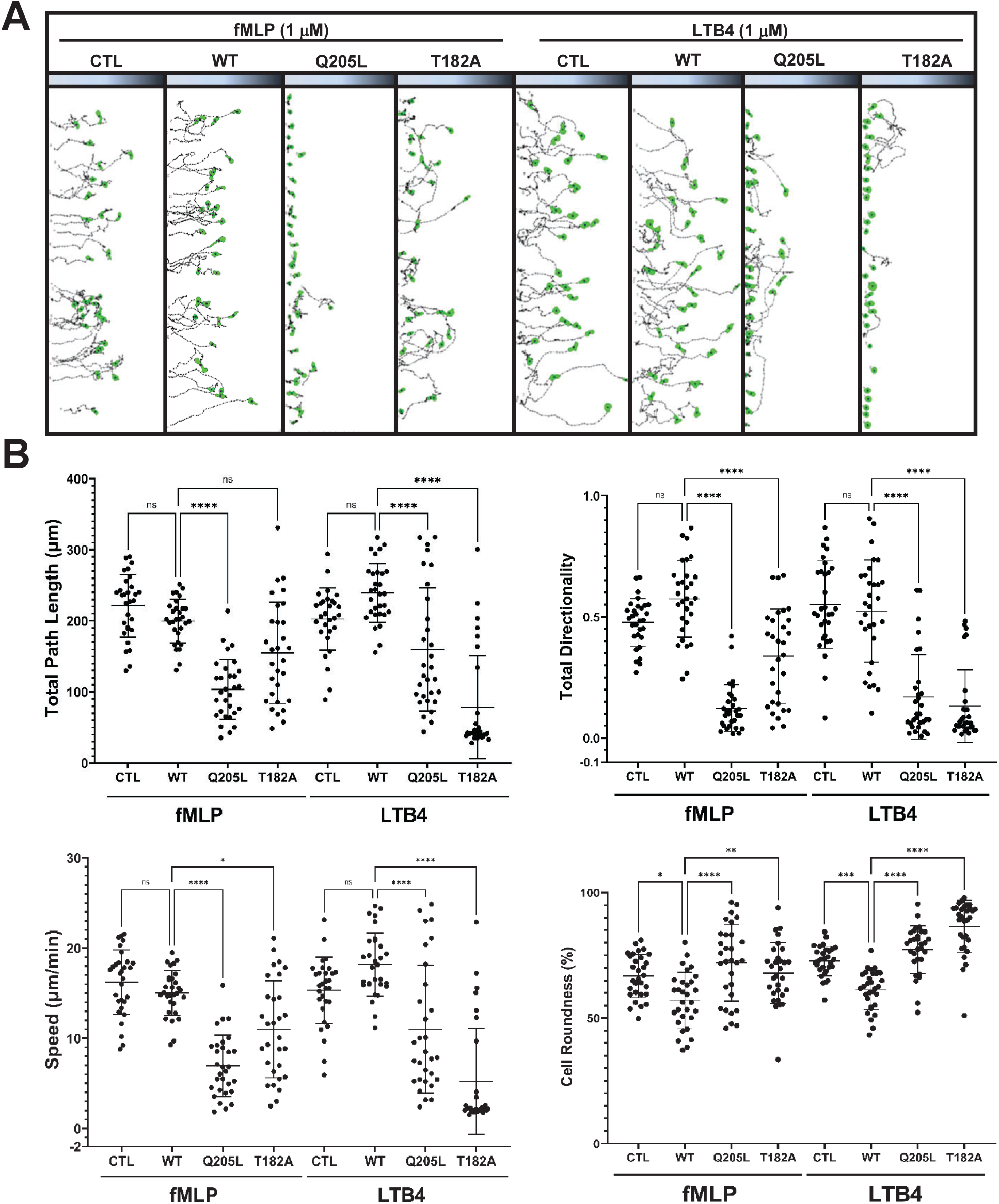
Impaired chemotaxis of CA-Gαi2 expressing neutrophils in the gradients at saturating chemoattractant gradients. **A.** Montages showing the travel path of chemotaxing neutrophils expressing WT, Q205Lor T182A of Gai2 in gradients generated from either saturating (1 mM) fMLP or LTB4 sources. The shaded panels on top of the images in the montage indicate the gradients. Gradients generated on the terrace are linear gradients. The concentration on the left side of the terrace is 0 and the concentration on the right side of the terrace is 1 mM. Movement of at least 30 cells in each group from multiple independent experiments under the same condition was analyzed by DIAS software. See **Video S5** for a complete set of cell migration. **B.** Chemotaxis behaviors measured from **A** are described as four parameters: directionality, speed, the total path length, and roundness as described in Materials and Methods. Thirty cells from each group were measured. Dot-plot analysis of the chemotaxis index was shown. Student’s t-test was used to calculate the *p* values, which are indicated as *ns* (not significant, ns, *p* > 0.05), * (*p* < 0.05), ** (*p* < 0.01), *** (*p* < 0.001), or **** (*p* < 0.0001).

### Increased Cell Sensitivity and Cell Migration in Neutrophils Expressing Active Gαi2 Mutants

Elevated Ras activity is known to enhance cellular sensitivity and migratory responses (Xu et al., 2021a; Xu et al., 2025; Xu et al., 2021b). To examine this effect in HL60 cells expressing constitutively active Gai2 mutants, we measured Ras activation following stimulation with a subsensitive concentration (0.1 nM) of fMLP. This concentration of fMLP induced minimally detectable Ras activation in WT-Gai2–expressing cells (**Figure 6A**) (Xu et al., 2021b). However, the same concentration of fMLP induced a clearly detectable, increased Ras activation in cells expressing either Q205L or T182A mutants of Gαi2 (**Figure 6A-6B**; **Figure S7**), indicating enhanced cellular sensitivity to chemoattractant stimulation.

**Figure 6.**
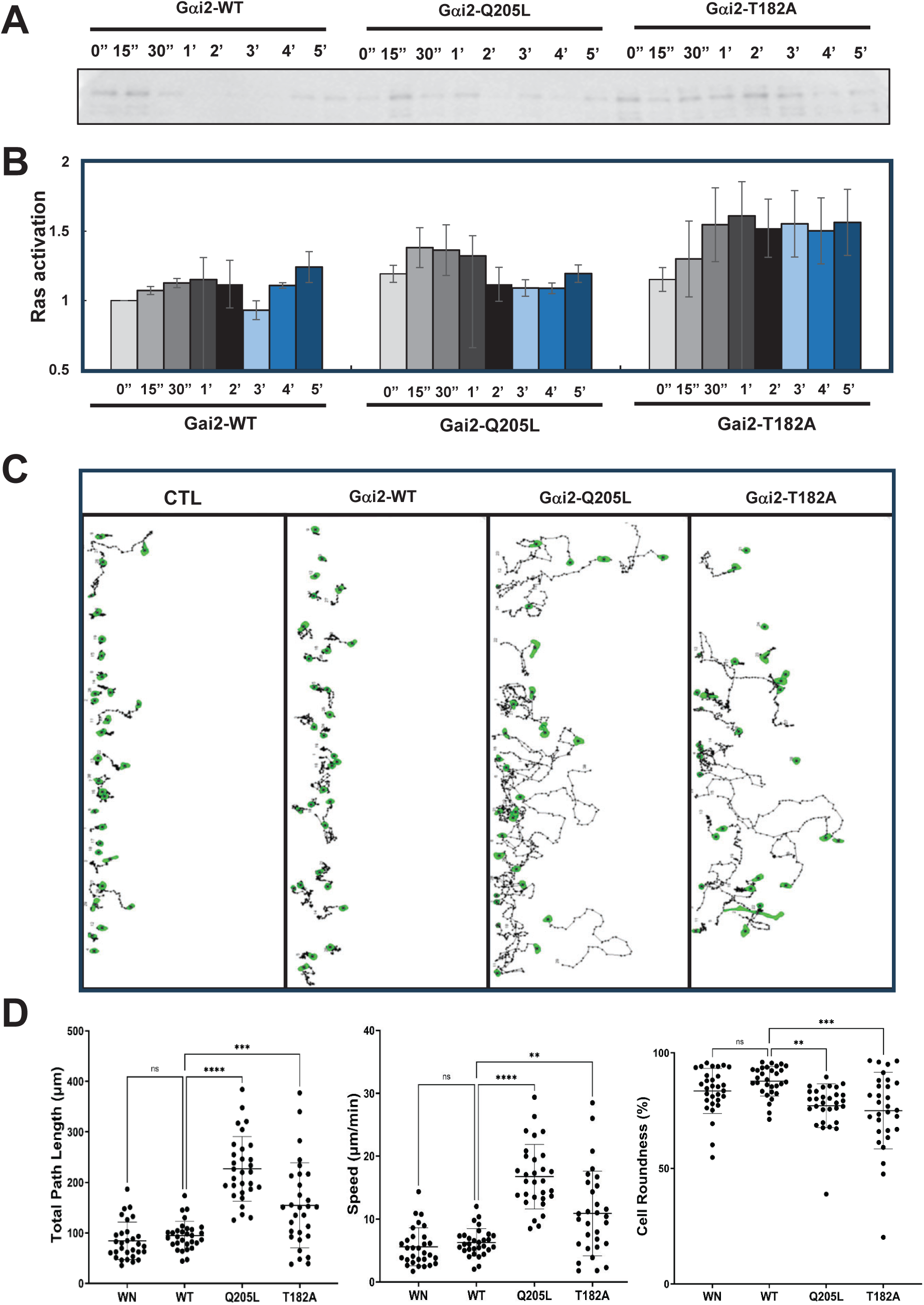
Increased cell sensitivity and cell motility of neutrophils expressing CA mutants of Gαi2. **A.** Ras activation in neutrophils expressing WT, Q205L, or T182A of Gai2 in response to subsensitive (0.1 nM) fMLP stimulation, as determined by pull-down assays. Cells were stimulated with fMLP at a final concentration of 0.1 nM at time 0s. Aliquots of cells were taken before or after fMLP at the indicated time points and subjected to the pull-down assay and subsequent western blot detection of active Ras as previously described. **B.** Quantitative measurement of Ras activation in the cells expressing either WT or CA mutants of Gai2 in response to fMLP stimulation in three independent experiments, including the result presented in **A** and two sets of independent measurements (**Figure S8**). The intensity ratio of active Ras in WT at time 0 s was normalized to 1. Mean ± SD from the three independent experiments is shown. **C.** Montages showing the random traveling path of neutrophils expressing endogenous Gai2 (CTL), or overexpressing WT or Q205L or T182A mutans of Gai2. Movement of 30 cells in each group was analyzed by DIAS software and is shown. For a full set of cell migration in each group of cells, see **Video S6**. **D.** Moving behaviors measured from **A** are described by three parameters: speed, total path length, and roundness as described in Figure 5B. Student’s t-test was used to calculate the *p* values, which are indicated as ns (not significant p > 0.05), * (p < 0.05), ** (p < 0.01), *** (p < 0.001), or **** (p < 0.0001).

Active Ras signaling is a hallmark of cell motility in both the model organism *Dictyostelium discoideum* and mammalian neutrophils. To determine whether increased Ras activity influences cell migration, we analyzed the random migratory behavior of HL60 cells expressing endogenous Gai2 or overexpressing WT-, Q205L-, or T182A-Gai2 (**Figure 6C-6D**; **Video S5**). Cells expressing endogenous or WT-Gai2 exhibited comparable migration behaviors. By contrast, cells expressing Q205L or T182A mutants displayed significantly enhanced random motility, with increased cell speed, total travel distance, and cell polarity. These results indicated that constitutively active Gai2 mutants enhance cellular sensitivity to chemoattractant stimulation and promote motility, likely through sustained Ras activation.

### Active Gαi2-Expressing HL60 Cells Exhibit Normal chemotaxis at Low Concentrations but Enhanced Chemotaxis at Subsensitive Concentrations

We previously showed that *capri* knockdown (*capri^kd^*) neutrophils exhibit increased chemoattractant sensitivity accompanied by elevated activation of Ras and its downstream effectors (Xu et al., 2021b). Notably, *capri^kd^* neutrophils display an altered chemotactic profile characterized by enhanced chemotaxis in gradients at subsensitive concentrations, normal chemotaxis at intermediate concentrations, and impaired chemotaxis at saturating concentrations, indicating a concentration-range upshift in chemotactic competency (Xu and Jin, 2022). Consistent with this phenotype, neutrophils expressing Gai2-Q205L or Gai2-T182A mutants exhibit impaired chemotaxis in gradients of saturating chemoattractants (**Figure 5**). To further investigate this behavior, we analyzed the chemotactic responses of neutrophils expressing WT-, Q205L-, or T182A-Gai2 in fMLP gradients at either intermediate (10 nM) or subsensitive (0.1 nM) concentrations (**Figure 7A–7B**; **Video S6**). In intermediate (10 nM) fMLP gradients, all groups displayed comparable chemotactic responses. However, in subsensitive (0.1 nM) fMLP gradients, most WT-Gai2 cells exhibited either random migration or minimal directional movement, whereas cells expressing Q205L or T182A mutants showed clear directional chemotaxis toward the gradient source. A similar pattern was observed in LTB4 gradients, where mutant-expressing cells demonstrated enhanced chemotaxis at subsensitive concentrations (0.1 nM) but displayed responses comparable to WT cells at intermediate concentrations (100 nM) (**Figure 7C–7D**; **Video S7**). Taken together, these results indicate that Q205L- and T182A-Gai2–expressing neutrophils exhibit concentration-dependent chemotactic behavior, characterized by enhanced chemotaxis in subsensitive gradients and normal responses at intermediate concentrations. Combined with the results from **Figures 5** and **6**, these findings demonstrate that constitutively active Gai2 mutants shift the functional range of neutrophil chemotaxis toward higher chemoattractant concentrations, enabling effective responses across an upshifted concentration window.

**Figure 7.**
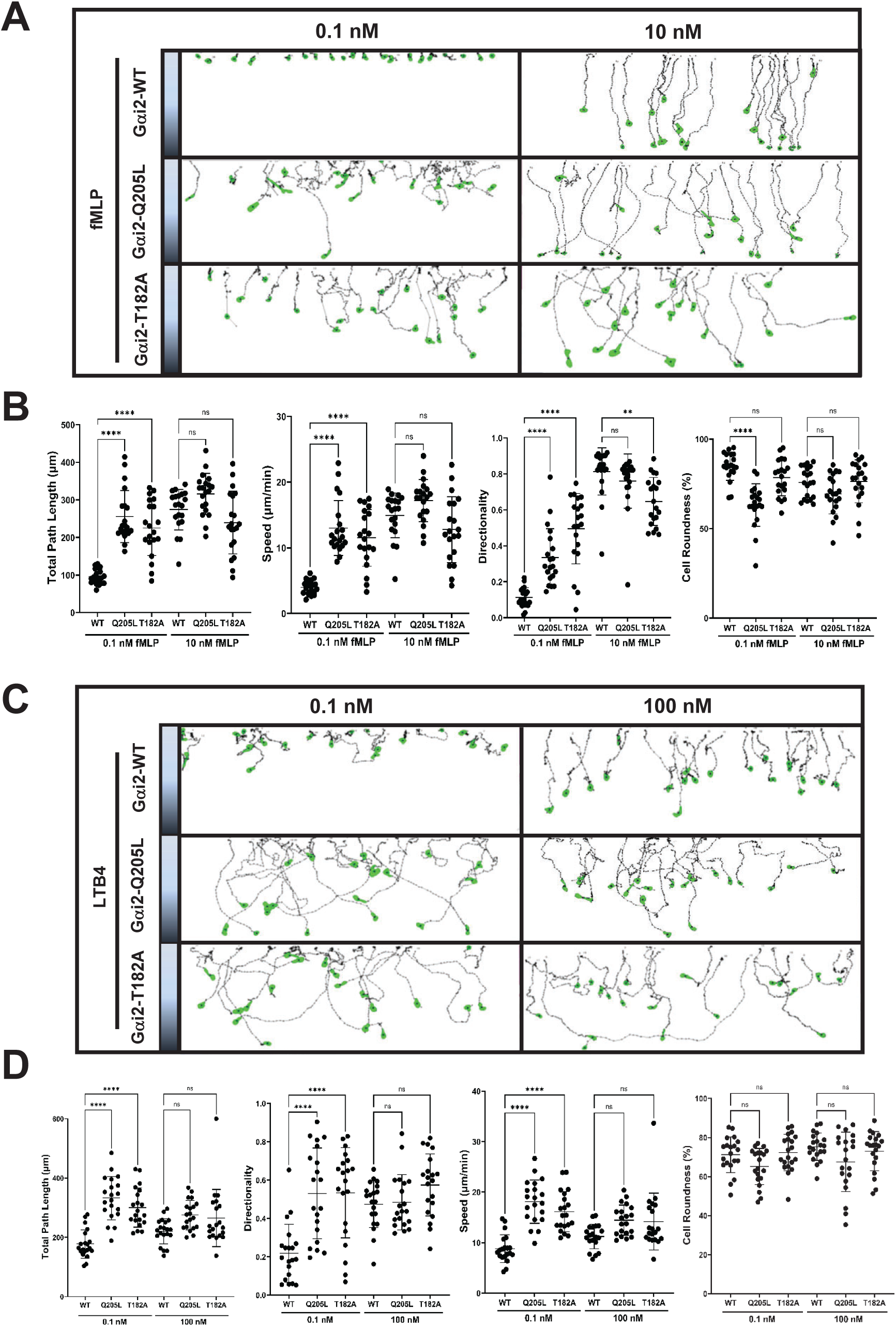
CA-Gαi2-expressing neutrophils display normal chemotaxis in the chemoattractant gradients at intermediate concentrations, but exhibit enhanced chemotaxis in gradients at subsensitive concentrations. **A.** Montages show the travel paths of neutrophils expressing WT or CA-mutans of Gai2 in response to gradients generated from source at the concentrations of 0.1 nM and 10 nM fMLP. The shaded panels on the left side of the images in the montage indicate gradient sourced from chemoattractants at the indicated concentrations. The concentration on the top side of the terrace is 0 and the concentration at the bottom side of the terrace is as indicated on the left side of the terrace. Movement of at least 30 cells in each group was analyzed by DIAS software and is shown. For complete set of cell migration, see **Video S7** (fMLP). **B.** Chemotaxis behaviors measured from **A** are described by four parameters: speed, total path length, and roundness. Student’s t-test was used to calculate the p values, which are indicated as ns (not significant *p* > 0.05), * (*p* < 0.05), ** (*p* < 0.01), *** (*p* < 0.001), or **** (*p* < 0.0001). **C.** Montages show the travel paths of neutrophils expressing WT or CA-mutans of Gai2 in response to gradients generated from source at the concentrations of 0.1 nM and 10 nM LTB4. The shaded panels on the left side of the images in the montage indicate gradient sourced from chemoattractants at the indicated concentrations. The concentration on the top side of the terrace is 0 and the concentration at the bottom side of the terrace is as indicated on the left side of the terrace. Movement of at least 30 cells in each group was analyzed by DIAS software and is shown. For complete set of cell migration, see **Video S8**. **D.** Chemotaxis behaviors measured from **C** are described by four parameters: speed, total path length, and roundness. Student’s t-test was used to calculate the p values, which are indicated as ns (not significant *p* > 0.05), * (*p* < 0.05), ** (*p* < 0.01), *** (*p* < 0.001), or **** (*p* < 0.0001).

**Figure 8.**
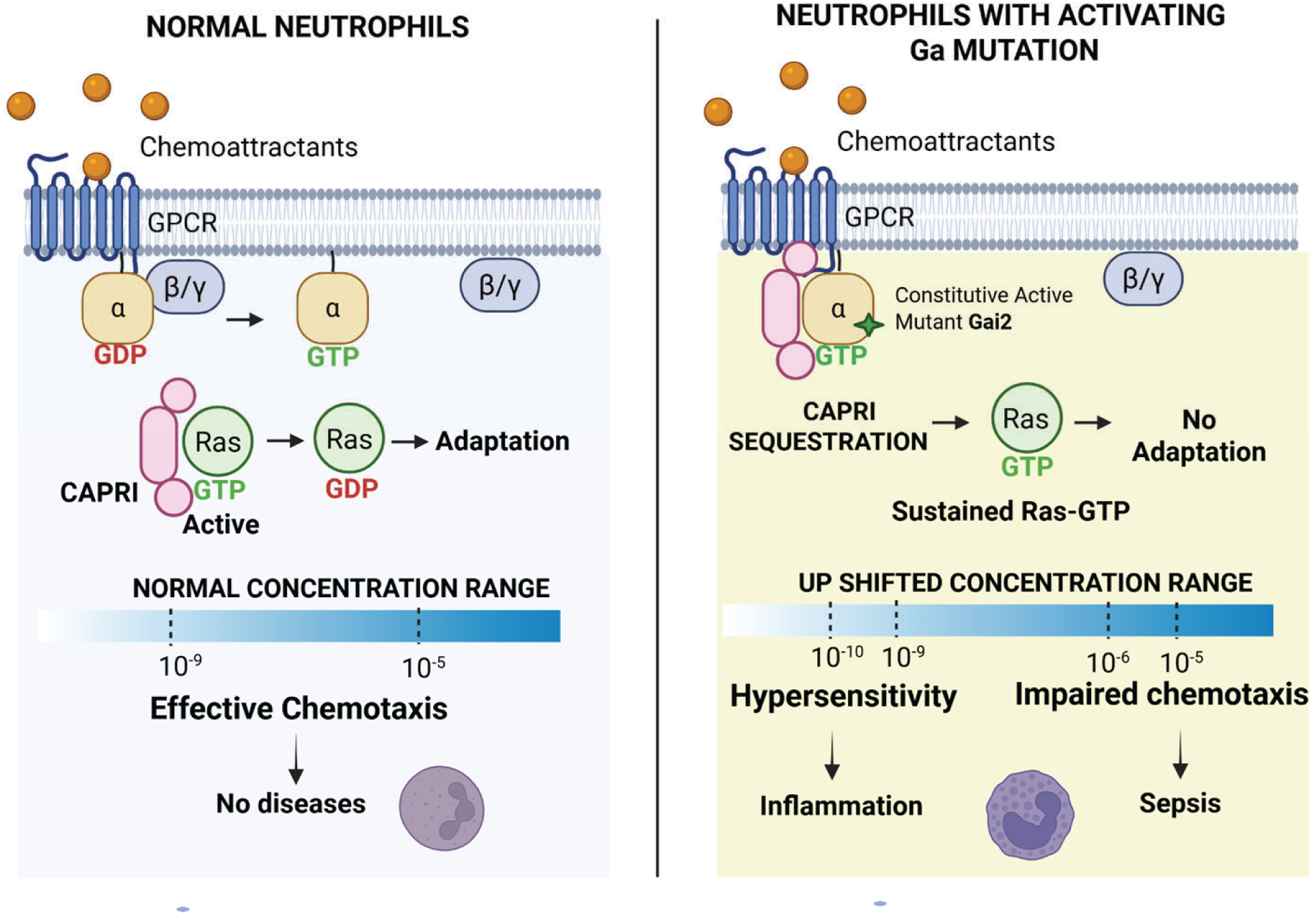
**CA-Gαi2 mutants binds CAPRI to upshift chemoattractant concentration range for chemotaxis.**

## DISCUSSION

Neutrophils sense chemoattractant gradients through G protein–coupled receptors (GPCRs) and heterotrimeric G proteins, which translate extracellular cues into intracellular signals that drive chemotaxis. Upon GPCR activation, G proteins dissociate into Ga and GBy subunits. Several direct effectors of GBy, including phospholipase C B2/3 (PLCB2/3), phosphoinositide 3-kinase y (PI3Ky), and P-Rex, mediate key signaling pathways controlling neutrophil migration (Kuang et al., 1996; Suire et al., 2006; Welch et al., 2002). In contrast, although Gai2 is the predominant Ga subunit in neutrophils and is essential for chemoattractant signaling, its downstream effectors remain poorly defined. Recently, interactions between RasGAP, RASA2, and Gai2, including both wild-type (WT) and activating mutants, have attracted attention due to their impact on immune signaling (Ham et al., 2024b), providing a potential link between Gai and Ras pathways beyond canonical GPCR activation. Here, we show that CAPRI directly associates with multiple Ga subunits, with constitutively active Gai2 exhibiting enhanced interaction. Notably, CAPRI interacted with multiple Gai subunits (**Figure 1C**). And, Gai2 also interacted with several RasGAPs, including RASA2, CAPRI, and RASAL3 (a RasGAP that is the second most abundant in neutrophils and contributes to neutrophil chemotaxis) (Lin et al., 2024) (**Figure 1D**). Consistent with the above, in the model organism of *D. discoideum*, the heterotrimeric G protein α2 subunit (Ga2) interacts with C2GAP1, a key component of gradient sensing to mediates Ras adaptation and chemotaxis (Xu et al., 2026). Together, these findings suggest that interactions between Ga subunits and RasGAPs may represent an evolutionarily conserved mechanism coordinating heterotrimeric G protein and Ras signaling. More importantly, cells lacking C2GAP1 (*c2gapA^−^*) exhibit enhanced heterotrimeric G protein activation upon chemoattractant stimulation, suggesting that this interaction negatively regulates G protein signaling. In the present study, our results reveal the functional impact of Gai–CAPRI interaction on RasGAP signaling and chemotaxis, although its role in regulating heterotrimeric G protein dynamics remains to be determined.

One remarkable feature of eukaryotic chemotaxis is the ability of cells to sense and respond to chemoattractant gradients spanning a broad concentration range. For example, cells detect approximately 10-s to 10-9 M cAMP in *Dictyostelium discoideum* and a similar range (10-s–10-9 M) of fMLP or SDF-1a in mammalian neutrophils. Ras is a central regulator of eukaryotic chemotaxis and has been extensively studied in both systems. In *D. discoideum*, loss of the Ras GTPase–activating protein C2GAP1 (c2gapA-) results in elevated basal Ras activity, defective Ras adaptation, and impaired chemotaxis under saturating stimulation. Notably, although these cells migrate poorly in high chemoattractant concentrations, they exhibit normal chemotaxis in intermediate gradients and enhanced sensitivity with improved migration at low or subsensitive concentrations (Xu et al., 2021a; Xu et al., 2017). Consequently, the effective chemotactic concentration range shifts downward from ∼10-s–10-9 M in wild-type cells to ∼10-6–10-¹0 M in *c2gapA-* cells. A similar regulatory mechanism operates in mammalian neutrophils. CAPRI-deficient (*capri^kd^*) human neutrophils display a comparable downward shift in chemotactic sensitivity across multiple chemoattractants (Xu et al., 2021b). CAPRI membrane recruitment depends on calcium signaling mediated by PLCy2 (Xu et al., 2023), and PLCy2 knockdown neutrophils phenocopy this behavior, showing impaired chemotaxis in saturating gradients but preserved or enhanced migration in intermediate or subsensitive gradients (Xu et al., 2025). Here, we further establish a conserved paradigm in which hyperactivation of Ras signaling redefines the chemotactic dynamic range. Neutrophils expressing constitutively active Gai2 display elevated basal Ras activity and migrate efficiently at low or subsensitive chemoattractant concentrations (**Figures 6** and **7**). Consistently, mouse neutrophils harboring constitutively active Gai2 (Gai2-G184S/G184S) exhibit increased chemotactic sensitivity (Cho et al., 2012), and T cells with defective RASA2 show heightened antigen responsiveness (Carnevale et al., 2022). Together, our findings demonstrate that the Gai2–CAPRI–Ras signaling axis functions as a dynamic range–tuning module in leukocytes. Rather than merely attenuating signaling amplitude, RasGAP-mediated regulation expands the operational window of chemotaxis, enabling cells to balance sensitivity to weak cues with robustness under strong stimulation. Dysregulation of this axis compresses the functional range, biasing cells toward heightened low- dose sensitivity at the expense of performance in high-concentration environments. Altogether, this new conceptual framework not only advances our mechanistic understanding of immune cell navigation but also identifies the Gai2–CAPRI–Ras node as a potential therapeutic target for modulating leukocyte responsiveness during inflammatory disorders and sepsis.

## ACKNOWLEDGMENTS

This research was supported by the Intramural Research Program of the National Institutes of Health (NIH). The contributions of the NIH author(s) were made as part of their official duties as NIH federal employees, are in compliance with agency policy requirements, and are considered Works of the United States Government. However, the findings and conclusions presented in this paper are those of the author(s) and do not necessarily reflect the views of the NIH or the U.S. Department of Health and Human Services.

## MATERIALS AND METHODS

### Cells, cell lines, cell culture, and differentiation

HL60 cells were cultured, maintained, and differentiated as previously reported (Xu et al., 2015). Briefly, cells were maintained in RPMI 1640 culture medium [RPMI 1640 medium with 10% (v/v) fetal bovine serum and 25 mM HEPES (Quality Biological, Inc. Gaithersburg, MD)]. Cells expressing endogenous Gαi2, or overexpressing WT, Q205L, or T182A mutant of Gαi2 were constructed as previously reported (Ham et al., 2024b). HL60 cells were differentiated in RPMI1640 culture medium containing 1.3% DMSO for 5 days before the experiments. The cells were incubated at 37°C in a humidified 5% CO_2_ atmosphere.

### Reagents and antibodies

IL8 and LBT4 were purchased from R&D Systems (Minneapolis, MN); f-Met-Leu-Phe (fMLP) and DMSO from Sigma Aldrich (St. Louis, MO); Anti-Gϕ3, -ϕ31, - ϕ32, -Gαi1/2, or -Gαi3 from Santa Cruz Biotechnology, Inc. (Dallas, TX); anti-CAPRI antibody from antibodies-online (Dunwoody, GA); anti-RASA2 rabbit antibodies from Biorbyt (Cambridge, UK); anti-pan Ras antibodies from EMD Millipore (Darmstadt, Germany); anti-actin and -CD11 from Cell signaling, HRP-conjugated goat anti-mouse and -rabbit antibodies, and HRP-conjugated donkey anti-goat IgG from Jackson ImmunoResearch (West Grove, PA); most tissue culture reagents from Life Technologies Inc. (Carlsbad, CA); and HEPES from Quality Biological, Inc. (Gaithersburg, MD).

### Immunoprecipitation assay

Briefly, HL60 cells expressing turbo-GFP (tGFP) alone or tGFP tagged WT/mutants of CAPRI (Figure 1C) or GFP-tagged Gαi2-WT, Gαi2-Q205L or Gαi2-T182A (Figure 1D) were starved with RPMI 1640 medium containing 25 mM HEPES at 37C for 3 hours as previously described. Cells were then collected and resuspended at 2×10^7^ cells/ml and then stimulated with a final concentration of 10 μM fMLP. Aliquots of cells at indicated time points were lysed with 10 ml immunoprecipitation buffer (IB, 20 mM Tris, pH 8.0; 20 mM MgCl_2_; 10 % glycerol; 2 mM Na_3_VO_4_; 0.25 % NP40; and complete 1X EDTA-free proteinase inhibitor) with or without 1 mM GTPγS for 30 min on ice. Cell extracts were centrifuged at 16,000 × *g* for 10 min at 4 C. Supernatant fractions were collected and incubated with IgG beads pre-conjugated with anti-GFP at 4 C for 2 hours. Beads were washed four times with immunoprecipitation buffer, and proteins were eluted by boiling the beads in 50 μl 2X sample loading buffer (SLD) (Quality Biological Inc, Gaithersburg, MD). The eluted samples were subjected to western blot detection of the indicated proteins.

### Ras activation assay

The procedure was as previously reported (Xu et al., 2021b). Cells were collected, resuspended at 2×10^7^ cells/ml, and transferred to a medical cup under constant shaking at 200 rpm for 3 min at room temperature. Cells were stimulated with fMLP at the indicated final concentrations. At the indicated time points before or after stimulation, equal number of cells were subject to the subsequent steps. Aliquots were then mixed with immunoprecipitation buffer (IB), including 0.25 % NP40, 10 mM Tris (pH7.5) buffer, 150 mM NaCl, 1 mM Na_3_VO_4_, 10 mM NaF, and 1X proteinase inhibitor (Roche, Basel, Switzerland). The mixtures were incubated on ice for 30 min and then centrifuged at 100,000 × *g* at 4 C for 30 min. The supernatants were incubated with agarose beads conjugated with RBD (active Ras binding domain of human Raf1) (Cytoskeleton, Inc. Denver, CO at 4 C for 2 hours. The agarose beads were washed three times with IB. The protein on the beads was eluted by mixing with 25 μl 2X sample loading buffer (SLD) (Quality Biological Inc, Gaithersburg, MD). The supernatants and eluted proteins were subjected to western blot detection of the indicated proteins.

### Immunoblotting

The detection of heterotrimeric G protein subunits in **Figure 1A, 1B**, and **1C** was carried out with membranes obtained previously (Xu et al., 2021b). The membranes with the proteins were incubated with the indicated antibodies by following the manufacturer’s instructions and washed three times with 10 μM Tris Buffer with 15 μM saline and 0.02 % Tween 20 (TBS-T) (Sigma Aldrich, St. Louis, MO) for 10 min. The membranes were incubated with the secondary antibodies by following the manufacturer’s instructions, washed three times with TBS-T for 10 min and subjected to detection of the indicated protein using the substrates of HRP (Life Technology, Carlsbad, CA).

### AlphaFold3-assisted analysis of Gα2-C2GAP1 interaction using NIH Biowulf cluster

Protein–protein complex models were generated using AlphaFold3 (DeepMind, 2024) installed as singularity container with af3 wrapper script (https://github.com/NIH-HPC/af3) on the NIH Biowulf high-performance computing cluster. Model parameters were downloaded from Google DeepMind with permission for non-commercial use. Protein sequences were obtained from uniprot.org. Jobs were submitted using the default AlphaFold3 (AF3) settings. Predictions were generated for human Gαi2 (-GDP or -GTP bound), CAPRI, Gϕ31, Gϕ32, and Gγ1 along with six interactions, including Gαi2Gϕ31, Gαi2Gϕ32, Gαi2Gϕ32Gγ(-GDP or -GTP bound), and CAPRI-Gα2 (-GDP or -GTP bound) of WT, Q205L, or T182A mutants. Five predictions were generated for each job with the top-ranked structure saved in CIF and JSON format and then accessed and visualized in ChimeraX version 1.11.1. Model accuracy was annotated in ChimeraX using the AlphaFold palette, with color-scheme mapped to per-residue confidence scores (pLDDT) accessed from the B-factor field of each respective CIF file. The predicted structures and parameters shown were calculated using PRODIGY based on instructions from https://github.com/haddocking/prodigy (Xue et al., 2016). Predictions were saved in .PDB format in ChimeraX and edited internally for accurate designation of atom identity and chain assignment to enable compatibility with PRODIGY.

### Imaging and data processing

Cells were plated and allowed to adhere to the cover glass of a 4-well or a 1-well chamber (Nalge Nunc International, Naperville, IL) precoated with Fibronectin (Sigma Aldrich, Saint Louis, MO) for 10 min and then covered with RPMI 1640 medium with 10% FBS and 25 mM HEPES. For confocal microscopy, cells were imaged using a Carl Zeiss Laser Scanning Microscope Zen 780 (Carl Zeiss, Thornwood, NY) with a Plan-Apochromat 40x/1.4 Oil DIC M27 objective. For the uniform-stimulation experiment of membrane translocation assays, the stimuli were directly delivered to the cells as previously described (Xu et al., 2016). Membrane translocation of the indicated protein was measured by the depletion of the interested protein in the cytoplasm as previously described (Xu et al., 2021b). For quantitative analysis of membrane translocation dynamics of the indicated molecules, the cytosolic depletion of the indicated molecule was measured. Regions of interest (ROIs) in the cytoplasm (avoiding the nucleus area as much as possible) remained within the cells throughout the time period of the measurements. To normalize the effect of photobleaching and morphological changes during data acquisition, the intensity of ROIs in the cytoplasm was divided by the intensity of whole cells at each given time point. Lastly, the resulting data were divided by that at time 0 s; consequently, the relative intensity of any cells at time 0 s became 1. The graph of mean ± SD is shown.

### TAXIScan chemotaxis assay and data analysis

The procedure was as previously reported (Wen et al., 2016). Briefly, differentiated cells were loaded onto one side of fibronectin-coated 4-μm EZ-TAXIScan chambers. The chemoattractants at the indicated concentrations were added to the other side of the well across the terrace to generate a liner gradient in which the cells migrated through. The traveled distance is the traveled length (μm). The length of terrace is the total length of the gradient generated (260 μm). The chemoattractant concentration of gradient (C) cells experienced at the each give position depends on both the concentration of fMLP source (C_source_) and the distance they traveled (the traveled length) from no gradient to the total distance (the total length) to the fMLP source. That is, C = C_source_ × ratio of the traveled length vs the total length). The cells migrated for 30 min at 37 °C. Images were taken for 30 min at 15 s intervals. For chemotaxis parameter measurements, 20 or 30 cells in each group were analyzed with DIAS software (Wessels et al., 1998) in Figure 5 and 7, respectively. Chemotaxis behaviors measured are described as four parameters: directionality, which is “upward” directionality, where 0 represents random movement and 1 represents straight movement toward the gradient; speed, defined as the distance that the centroid of the cell moves as a function of time; total path length, the total distance the cell has traveled; and roundness (%) for polarization, which is calculated as the ratio of the width to the length of the cell. Thus, a circle (no polarization) is 1 and a line (perfect polarization) is 0. The extracted data were analyzed by Prism and Student’s *t*-test was used to calculate the *p* values.

## ACKNOWLEDGMENTS

This research was supported by the Intramural Research Program of the National Institutes of Health (NIH). The contributions of the NIH author(s) are considered Works of the United States Government. The findings and conclusions presented in this paper are those of the author(s) and do not necessarily reflect the views of the NIH or the U.S. Department of Health and Human Services.

## CONFLICTS OF INTEREST

The authors declare that they have no conflict of interest.

## AUTHOR CONTRIBUTIONS

Conceptualization: X.X.; Investigation: X. X., W.K., R.K.; Data analysis: X. X., W.K., A.L., R.K., C.Z.; Writing – Original draft: X.X.; Review & Editing: X. X., W.K., A.L., R.K., C.Z., H.J., H.C.S., T.J.; Funding: T.J.

## Notes

### Competing Interest Statement

The authors have declared no competing interest.

## REFERENCES

Birnie, G.D., 1988. The HL60 cell line: a model system for studying human myeloid cell differentiation. Br J Cancer Suppl 9, 41–45.

Carnevale, J., Shifrut, E., Kale, N., Nyberg, W.A., Blaeschke, F., Chen, Y.Y., Li, Z., Bapat, S.P., Diolaiti, M.E., O’Leary, P., Vedova, S., Belk, J., Daniel, B., Roth, T.L., Bachl, S., Anido, A.A., Prinzing, B., Ibañez-Vega, J., Lange, S., Haydar, D., Luetke-Eversloh, M., Born-Bony, M., Hegde, B., Kogan, S., Feuchtinger, T., Okada, H., Satpathy, A.T., Shannon, K., Gottschalk, S., Eyquem, J., Krenciute, G., Ashworth, A., Marson, A., 2022. RASA2 ablation in T cells boosts antigen sensitivity and long-term function. Nature 609, 174–182.

Cho, H., Kamenyeva, O., Yung, S., Gao, J.-L., Hwang, I.-Y., Park, C., Murphy, P.M., Neubig, R.R., Kehrl, J.H., 2012. The Loss of RGS Protein-Gai2 Interactions Results in Markedly Impaired Mouse Neutrophil Trafficking to Inflammatory Sites. Molecular and Cellular Biology 32, 4561–4571.

Coleman, D.E., Berghuis, A.M., Lee, E., Linder, M.E., Gilman, A.G., Sprang, S.R., 1994. Structures of Active Conformations of Gi&#x3b1;1 and the Mechanism of GTP Hydrolysis. Science 265, 1405–1412.

Dessauer, C.W., Tesmer, J.J., Sprang, S.R., Gilman, A.G., 1998. Identification of a Gialpha binding site on type V adenylyl cyclase. J Biol Chem 273, 25831–25839.

Ham, H., Jing, H., Lamborn, I.T., Kober, M.M., Koval, A., Berchiche, Y.A., Anderson, D.E., Druey, K.M., Mandl, J.N., Isidor, B., Ferreira, C.R., Freeman, A.F., Ganesan, S., Karsak, M., Mustillo, P.J., Teo, J., Zolkipli-Cunningham, Z., Chatron, N., Lecoquierre, F., Oler, A.J., Schmid, J.P., Kuhns, D.B., Xu, X., Hauck, F., Al-Herz, W., Wagner, M., Terhal, P.A., Muurinen, M., Barlogis, V., Cruz, P., Danielson, J., Stewart, H., Loid, P., Rading, S., Keren, B., Pfundt, R., Zarember, K.A., Vill, K., Potocki, L., Olivier, K.N., Lesca, G., Faivre, L., Wong, M., Puel, A., Chou, J., Tusseau, M., Moutsopoulos, N.M., Matthews, H.F., Simons, C., Taft, R.J., Soldatos, A., Masle-Farquhar, E., Pittaluga, S., Brink, R., Fink, D.L., Kong, H.H., Kabat, J., Kim, W.S., Bierhals, T., Meguro, K., Hsu, A.P., Gu, J., Stoddard, J., Banos-Pinero, B., Slack, M., Trivellin, G., Mazel, B., Soomann, M., Li, S., Watts, V.J., Stratakis, C.A., Rodriguez-Quevedo, M.F., Bruel, A.-L., Lipsanen-Nyman, M., Saultier, P., Jain, R., Lehalle, D., Torres, D., Sullivan, K.E., Barbarot, S., Neu, A., Duffourd, Y., Similuk, M., McWalter, K., Blanc, P., Bézieau, S., Jin, T., Geha, R.S., Casanova, J.-L., Makitie, O.M., Kubisch, C., Edery, P., Christodoulou, J., Germain, R.N., Goodnow, C.C., Sakmar, T.P., Billadeau, D.D., Küry, S., Katanaev, V.L., Zhang, Y., Lenardo, M.J., Su, H.C., 2024a. Germline mutations in a G protein identify signaling cross-talk in T cells. Science 385, eadd8947.

Ham, H., Jing, H., Lamborn, I.T., Kober, M.M., Koval, A., Berchiche, Y.A., Anderson, D.E., Druey, K.M., Mandl, J.N., Isidor, B., Ferreira, C.R., Freeman, A.F., Ganesan, S., Karsak, M., Mustillo, P.J., Teo, J., Zolkipli-Cunningham, Z., Chatron, N., Lecoquierre, F., Oler, A.J., Schmid, J.P., Kuhns, D.B., Xu, X., Hauck, F., Al-Herz, W., Wagner, M., Terhal, P.A., Muurinen, M., Barlogis, V., Cruz, P., Danielson, J., Stewart, H., Loid, P., Rading, S., Keren, B., Pfundt, R., Zarember, K.A., Vill, K., Potocki, L., Olivier, K.N., Lesca, G., Faivre, L., Wong, M., Puel, A., Chou, J., Tusseau, M., Moutsopoulos, N.M., Matthews, H.F., Simons, C., Taft, R.J., Soldatos, A., Masle-Farquhar, E., Pittaluga, S., Brink, R., Fink, D.L., Kong, H.H., Kabat, J., Kim, W.S., Bierhals, T., Meguro, K., Hsu, A.P., Gu, J., Stoddard, J., Banos-Pinero, B., Slack, M., Trivellin, G., Mazel, B., Soomann, M., Li, S., Watts, V.J., Stratakis, C.A., Rodriguez-Quevedo, M.F., Bruel, A.L., Lipsanen-Nyman, M., Saultier, P., Jain, R., Lehalle, D., Torres, D., Sullivan, K.E., Barbarot, S., Neu, A., Duffourd, Y., Similuk, M., McWalter, K., Blanc, P., Bézieau, S., Jin, T., Geha, R.S., Casanova, J.L., Makitie, O.M., Kubisch, C., Edery, P., Christodoulou, J., Germain, R.N., Goodnow, C.C., Sakmar, T.P., Billadeau, D.D., Küry, S., Katanaev, V.L., Zhang, Y., Lenardo, M.J., Su, H.C., 2024b. Germline mutations in a G protein identify signaling cross-talk in T cells. Science 385, eadd8947.

Hewitt, N., Ma, N., Arang, N., Martin, S.A., Prakash, A., DiBerto, J.F., Knight, K.M., Ghosh, S., Olsen, R.H.J., Roth, B.L., Gutkind, J.S., Vaidehi, N., Campbell, S.L., Dohlman, H.G., 2023. Catalytic site mutations confer multiple states of G protein activation. Science Signaling 16, eabq7842.

Houk, A.R., Jilkine, A., Mejean, C.O., Boltyanskiy, R., Dufresne, E.R., Angenent, S.B., Altschuler, S.J., Wu, L.F., Weiner, O.D., 2012. Membrane tension maintains cell polarity by confining signals to the leading edge during neutrophil migration. Cell 148, 175–188.

Kuang, Y., Wu, Y., Smrcka, A., Jiang, H., Wu, D., 1996. Identification of a phospholipase C beta2 region that interacts with Gbeta-gamma. Proc Natl Acad Sci U S A 93, 2964–2968.

Kuwano, Y., Adler, M., Zhang, H., Groisman, A., Ley, K., 2016. Gai2 and Gai3 Differentially Regulate Arrest from Flow and Chemotaxis in Mouse Neutrophils. J Immunol 196, 3828–3833.

Lambright, D.G., Noel, J.P., Hamm, H.E., Sigler, P.B., 1994. Structural determinants for activation of the alpha-subunit of a heterotrimeric G protein. Nature 369, 621–628.

Lambright, D.G., Sondek, J., Bohm, A., Skiba, N.P., Hamm, H.E., Sigler, P.B., 1996. The 2.0 Å crystal structure of a heterotrimeric G protein. Nature 379, 311–319.

Lin, Y., Pal, D.S., Banerjee, P., Banerjee, T., Qin, G., Deng, Y., Borleis, J., Iglesias, P.A., Devreotes, P.N., 2024. Ras suppression potentiates rear actomyosin contractility-driven cell polarization and migration. Nat Cell Biol 26, 1062–1076.

Lockyer, P.J., Kupzig, S., Cullen, P.J., 2001. CAPRI regulates Ca(2+)-dependent inactivation of the Ras-MAPK pathway. Curr Biol 11, 981–986.

Masuho, I., Balaji, S., Muntean, B.S., Skamangas, N.K., Chavali, S., Tesmer, J.J.G., Babu, M.M., Martemyanov, K.A., 2020. A Global Map of G Protein Signaling Regulation by RGS Proteins. Cell 183, 503–521.e519.

Neves, S.R., Ram, P.T., Iyengar, R., 2002. G Protein Pathways. Science 296, 1636–1639.

Rasmussen, S.G., DeVree, B.T., Zou, Y., Kruse, A.C., Chung, K.Y., Kobilka, T.S., Thian, F.S., Chae, P.S., Pardon, E., Calinski, D., Mathiesen, J.M., Shah, S.T., Lyons, J.A., Caffrey, M., Gellman, S.H., Steyaert, J., Skiniotis, G., Weis, W.I., Sunahara, R.K., Kobilka, B.K., 2011. Crystal structure of the beta2 adrenergic receptor-Gs protein complex. Nature 477, 549–555.

Rincon, E., Rocha-Gregg, B.L., Collins, S.R., 2018. A map of gene expression in neutrophil-like cell lines. BMC Genomics 19, 573.

Suire, S., Condliffe, A.M., Ferguson, G.J., Ellson, C.D., Guillou, H., Davidson, K., Welch, H., Coadwell, J., Turner, M., Chilvers, E.R., Hawkins, P.T., Stephens, L., 2006. Gbetagammas and the Ras binding domain of p110gamma are both important regulators of PI(3)Kgamma signalling in neutrophils. Nat Cell Biol 8, 1303–1309.

Taussig, R., Iñiguez-Lluhi, J.A., Gilman, A.G., 1993. Inhibition of adenylyl cyclase by Gi alpha. Science 261, 218–221.

Wall, M.A., Coleman, D.E., Lee, E., Iñiguez-Lluhi, J.A., Posner, B.A., Gilman, A.G., Sprang, S.R., 1995. The structure of the G protein heterotrimer Gi alpha 1 beta 1 gamma 2. Cell 83, 1047–1058.

Welch, H.C., Coadwell, W.J., Ellson, C.D., Ferguson, G.J., Andrews, S.R., Erdjument-Bromage, H., Tempst, P., Hawkins, P.T., Stephens, L.R., 2002. P-Rex1, a PtdIns(3,4,5)P3- and Gbetagamma-regulated guanine-nucleotide exchange factor for Rac. Cell 108, 809–821.

Wen, X., Jin, T., Xu, X., 2016. Imaging G Protein-coupled Receptor-mediated Chemotaxis and its Signaling Events in Neutrophil-like HL60 Cells. J Vis Exp.

Wessels, D., Voss, E., Von Bergen, N., Burns, R., Stites, J., Soll, D.R., 1998. A computer-assisted system for reconstructing and interpreting the dynamic three-dimensional relationships of the outer surface, nucleus and pseudopods of crawling cells. Cell Motil Cytoskeleton 41, 225–246.

Xu, X., Bhimani, S., Pots, H., Wen, X., Jeon, T.J., Kortholt, A., Jin, T., 2021a. Membrane Targeting of C2GAP1 Enables Dictyostelium discoideum to Sense Chemoattractant Gradient at a Higher Concentration Range. Front Cell Dev Biol 9, 725073.

Xu, X., Gera, N., Li, H., Yun, M., Zhang, L., Wang, Y., Wang, J., Jin, T., 2015. GPCR-Mediated PLCbetagamma/PKCbeta/PKD Signaling Pathway Regulates the Cofilin Phosphatase Slingshot 2 in Neutrophil Chemotaxis. Molecular biology of the cell.

Xu, X., Jin, T., 2022. Ras inhibitors gate chemoattractant concentration range for chemotaxis through controlling GPCR-mediated adaptation and cell sensitivity. Front Immunol 13, 1020117.

Xu, X., Kim, R., Hyun, H., Shukla, R.d., Jin, T., 2026. Heterotrimeric G Protein and RasGAP Coupling Drives Adaptation During Chemotaxis. bioRxiv, 2026.2002.2024.707728.

Xu, X., Kim, W.S., Lee, A., Jin, T., 2025. PLCy2 controls neutrophil-like cell sensitivity through calcium oscillation and gates chemoattractant concentration range for chemotaxis. Frontiers in Immunology Volume 16–2025.

Xu, X., Wen, X., Bhimani, S., Moosa, A., Parsons, D., Ha, H., Jin, T., 2023. G protein-coupled receptor-mediated membrane targeting of PLCy2 is essential for neutrophil chemotaxis. J Leukoc Biol 114, 126–141.

Xu, X., Wen, X., Moosa, A., Bhimani, S., Jin, T., 2021b. Ras inhibitor CAPRI enables neutrophil-like cells to chemotax through a higher-concentration range of gradients. Proc Natl Acad Sci U S A 118.

Xu, X., Wen, X., Veltman, D.M., Keizer-Gunnink, I., Pots, H., Kortholt, A., Jin, T., 2017. GPCR-controlled membrane recruitment of negative regulator C2GAP1 locally inhibits Ras signaling for adaptation and long-range chemotaxis. Proc Natl Acad Sci U S A 114, E10092–e10101.

Xu, X., Yun, M., Wen, X., Brzostowski, J., Quan, W., Wang, Q.J., Jin, T., 2016. Quantitative Monitoring Spatiotemporal Activation of Ras and PKD1 Using Confocal Fluorescent Microscopy. Methods Mol Biol 1407, 307–323.

Xue, L.C., Rodrigues, J.P., Kastritis, P.L., Bonvin, A.M., Vangone, A., 2016. PRODIGY: a web server for predicting the binding affinity of protein–protein complexes. Bioinformatics 32, 3676–3678.

Yan, S.L., Hwang, I.Y., Kamenyeva, O., Kabat, J., Kim, J.S., Park, C., Kehrl, J.H., 2021. Unrestrained Ga(i2) Signaling Disrupts Neutrophil Trafficking, Aging, and Clearance. Front Immunol 12, 679856.

