## Supplementary material for "Heterotrimeric G protein αi2 Sequesters RasGAP to Control Neutrophil Sensitivity and Chemotaxis": SI

**Supplementary Information**

**Supplementary figure legends**

**Figure S1. Expression of wild-type (WT), or CA mutants (Q205L or T182A) of Gαi2 in HL60 cells.**

**A.** Montages show the detection of GFP-tagged WT, or CA mutants of Q205L or T182A in transduced or untransduced parent HL60 cells.

**B.** Quantification of the percentage of positive expression of WT or mutants of Gαi2 in HL60 cells.

**Figure S2. Expression of Ras GAPs in primary mammalian neutrophils and two leukocyte cell lines (HL60 and PLB985).** Data were extracted from https://pubmed.ncbi.nlm.nih.gov/30068296/.

**Figure S3. Predicted structures of the heterotrimeric Gαi2Gβγ.**

**A.** Predicted structures of Gαi2 (WT, Q205L, and T182A) in complex with GDP or GTP were generated using AlphaFold 3. Gαi2 comprises a Ras-like GTPase domain and an α-helical domain, separated by a deep cleft that accommodates the bound nucleotide. The N-terminus and C-terminus are indicated. In the GDP-bound state, GDP is depicted as red spheres, whereas GTP is shown as green spheres in the GTP-bound state. The point mutations Q205L and T182A are highlighted in cyan. The lower panel displays prediction confidence scores (pLDDT) from AlphaFold3, indicating the reliability of the modeled structures across residues.

**B.** Predicted complex of Gαi2Gβ1, Gαi2Gβ2, Gαi2Gβ2γ1), with Gα2 in either the GDP-(red, upper panel) or GTP-bound (green, lower panel) state. Gαi2 is shown in gold, Gβ in blue, and Gγ in dark magenta.

**Figure S4. Interaction interfaces and binding affinities between Gαi2 (WT or mutant) and CAPRI.**

**A.** Predicted complexes of CAPRI–Gαi2 interaction in which Gαi WT or mutant in GDP-bound state. WT, left panel; Q205L, middle panel; T182A, right panel. Gα2 (blue) in either GDP-bound (red spheres, top panels) or GTP-bound (green spheres, bottom panels) states. The point mutations Q205L and T182A are highlighted in magenta. The C2A (red), C2B (grey), GAP (gold), PH (purple), BTK (blue) and C-terminal (magenta) domains of CAPRI (cyan).

**B.** Predicted parameters of binding affinity between (GDP- or GTP-bound) Gα2 and C2GAP1 interaction.

**Figure S5.** **Ras activation in HL60 cells expressing WT, Q205L, or T182A mutant of Gαi2 upon saturating (1 μM) fMLP stimulation.** Two independent measurements are shown in **A** and **B** for the quantification of Ras activation in **Figure 4B**.

**Figure S6.** **Impaired chemotaxis of HL60 cells expressing either Gαi2-WT, -Q205L, or -T182A in the gradient of saturating (1 μM) SDF1a.**

**A.** Montages showing the travel path of neutrophils expressing WT or CA-mutans of Gαi2 in response to fMLP or LTB4 gradients generated from source of 1 μM SDF1α. The shaded panels on the left side of the images in the montage indicate gradient sourced from chemoattractants at the indicated concentrations. The concentration on the top side of the terrace is 0 and the concentration at the bottom side of the terrace is 1 μM. Movement of at least 30 cells in each group was analyzed by DIAS software and is shown.

**B.** Chemotaxis behaviors measured from **A** are described by four parameters: directionality, speed, total path length, and roundness as described in Materials and Methods. Student's t-test was used to calculate the p values, which are indicated as ns (not significant $p > 0.05$), * ($p < 0.05$), ** ($p < 0.01$), *** ($p < 0.001$), or **** ($p < 0.0001$).

**Figure S7.** **Ras activation in HL60 cells expressing Gαi2-WT, -Q205L, or -T182A upon subsensitive (0.1 nM) fMLP stimulation.** Two independent measurements of Ras activation are shown in **A** and **B** for the quantification shown in **Figure 6B**.

**Supplementary videos:**

**Video S1.** Montage showing alignment of Gαi2-WT in complexes in GDP-bound (grey; GDP, red spheres) and GTP-bound (gold; GTP, green spheres) states. Key residues exhibiting significant positional shifts in the switch region between the GDP- and GTP-bound forms are shown in the corresponding colors.

**Video S2** Montage showing alignment of Gαi2-WT (gold) and Gαi2-Q205L (grey) (top panel) and Gαi2-WT (gold) and Gαi2-T182A (grey) (lower panel) complexes in GDP-bound (left; GDP, red spheres) and GTP (right; GTP, green spheres) states. Point mutations of Q205L and T182A are highlighted in magenta. Key residues exhibiting significant positional shifts between the GDP- and GTP-bound forms are shown in the corresponding colors.

**Video S3.** Montage of CAPRI–Gαi2 (WT, Q205L, T182A) complexes in GDP-bound (top; GDP shown as red spheres) and GTP-bound (bottom; GTP shown as green spheres) states. Gαi2 is shown in light blue. CAPRI domains are color-coded as follows: C2A (red), C2B (grey), RasGAP (gold), PH (green), BTK (dark blue), and C-terminus (dark magenta). Point mutations of Q205L and T182A are highlighted in magenta.

**Video S4.** Monitoring Ras activation in HL-60 cells expressing GFP-tagged wild-type Gαi2 (Gαi2-WT-GFP, top panel), constitutively active Gαi2-Q205L (green, middle panel), or Gαi2-T182A (green, bottom panel) upon fMLP stimulation. Ras activation was assessed by plasma membrane (PM) translocation of the active Ras biosensor RBD-RFP. From left to right: Gαi2-WT or mutant (green), RBD-RFP (red), Differential interference contrast (DIC), and merged channels. Cells were stimulated with 1 µM fMLP at the beginning of the movies. Scale bar, 10 µm.

**Video S5.** Montage showing the migration of four types of HL60 cells: cells expressing endogenous Gαi2, or overexpressing Gαi2-WT, Gαi2-Q205L, or Gαi2-T182A (from top to bottom, respectively). Cells were exposed to gradients of 1 µM fMLP (left) or 1 µM LTB4 (right).

**Video S6.** Montage showing random migration of HL60 cells expressing endogenous wild-type Gαi2 (CTL), or overexpressing Gαi2-WT, Gαi2-Q205L (constitutively active), or Gαi2-T182A (constitutively active), from top to bottom, respectively.

**Video S7.** Montage showing the migration of HL60 cells expressing wild-type Gαi2 (top panels), Gαi2-Q205L (constitutively active, middle panels), or Gαi2-T182A (constitutively active, bottom panels) in gradients generated from localized sources of fMLP at subsensitive (0.1 nM, left) or intermediate (10 nM, right) concentrations.

**Video S8.** Montage showing the migration of HL60 cells expressing wild-type Gαi2 (WT), or the active mutants Gαi2-Q205L or Gαi2-T182A (top to bottom, respectively), in gradients generated from the sources of LTB4 at subsensitive (0.1 nM, left) or intermediate (100 nM, right) concentrations.

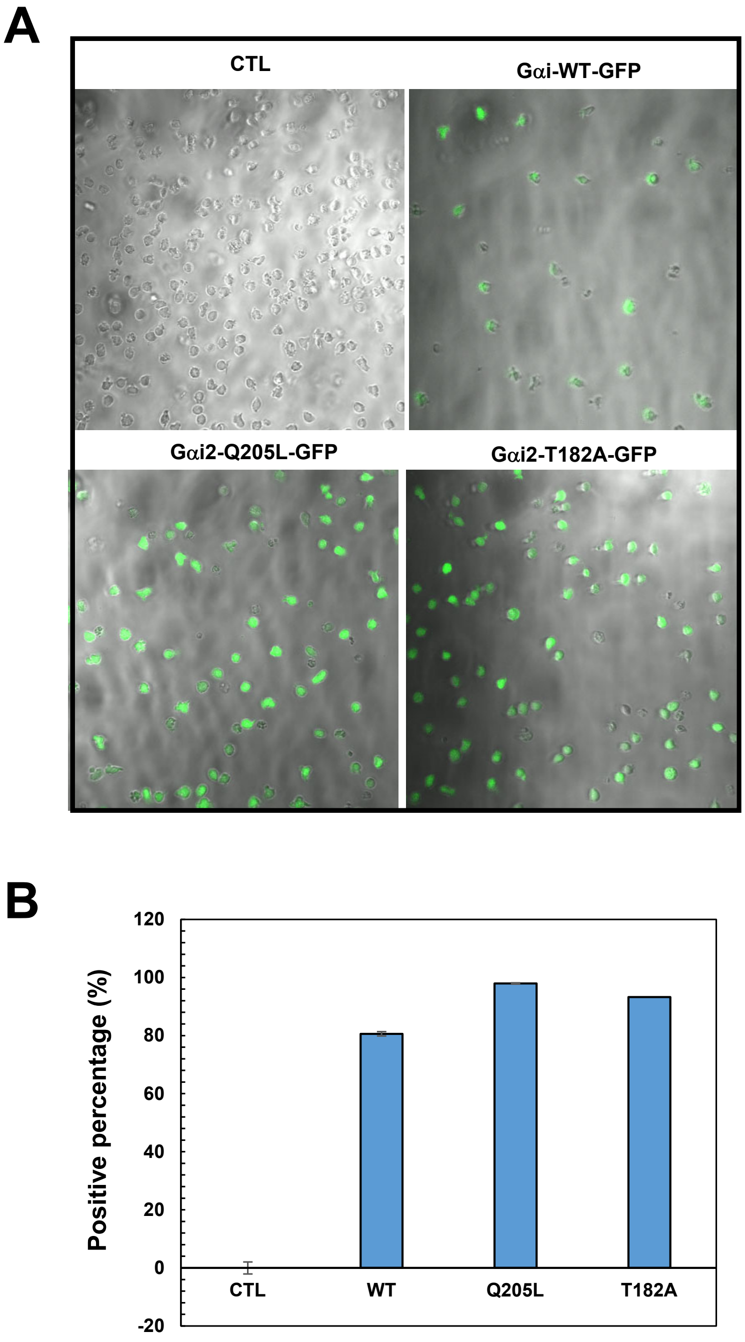
A
CTL
Gαi-WT-GFP
Gαi2-Q205L-GFP
Gαi2-T182A-GFP
B
Positive percentage (%)
120
100
80
60
40
20
0
-20
CTL
WT
Q205L
T182A

| | Hs | Mm | HL60 undiff | HL60 diff | PLB985 undiff | PLB985 diff |
|---|---|---|---|---|---|---|
| gap120 (RASA1) | 1.01 | 1.51 | 0.93 | 1.15 | 0.73 | 0.94 |
| RASA2 | 1.32 | 0.74 | 0.86 | 1.11 | 0.86 | 1.03 |
| CAPRI (RASA4) | 2.16 | 0.26 | -0.26 | 1.86 | -1.4 | 1.61 |
| RASAL1 | -2.39 | -0.48 | 0.85 | 1.38 | 0.87 | 1.05 |
| RASAL2 | -1.48 | -1.17 | 0.15 | -0.63 | 0.28 | -0.53 |
| RASAL3 | 1.54 | 1.51 | 0.93 | 1.15 | 0.73 | 0.94 |

Hs: Human primary neutrophils
Mm: Mice primary neutrophils
HL60 undiff: HL60 undifferentiated
HL60 diff: HL60 differentiated
PLB985 undiff: PLB985 undifferentiated
PLB985 diff: PLB985 differentiated
FPKM was normalized on a log10 scale.

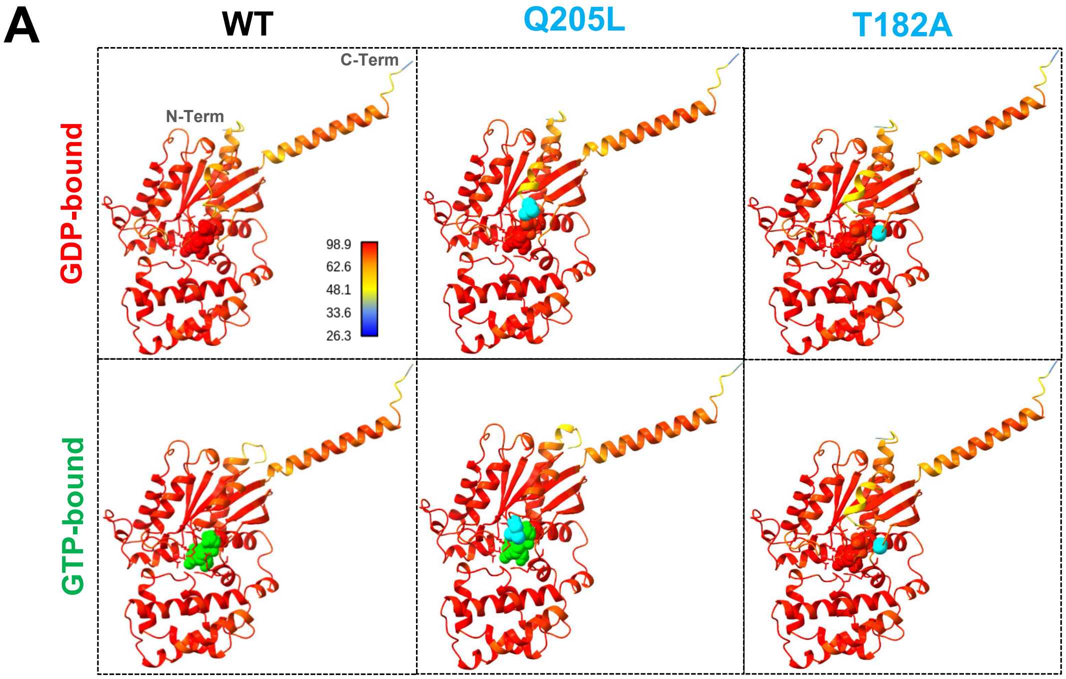
A
WT
Q205L
T182A
GDP-bound
GTP-bound
C-Term
N-Term
98.9
62.6
48.1
33.6
26.3

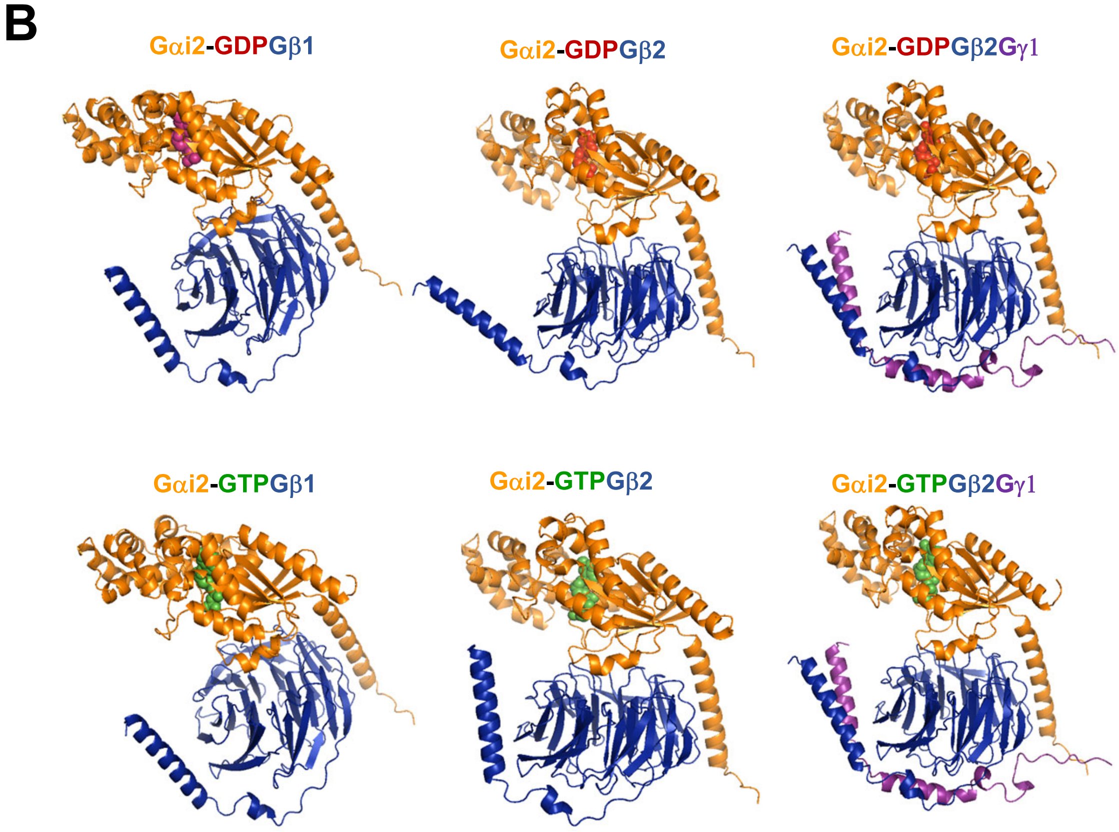
B
Gαi2-GDPGβ1
Gαi2-GDPGβ2
Gαi2-GDPGβ2Gγ1
Gαi2-GTPGβ1
Gαi2-GTPGβ2
Gαi2-GTPGβ2Gγ1

**A**

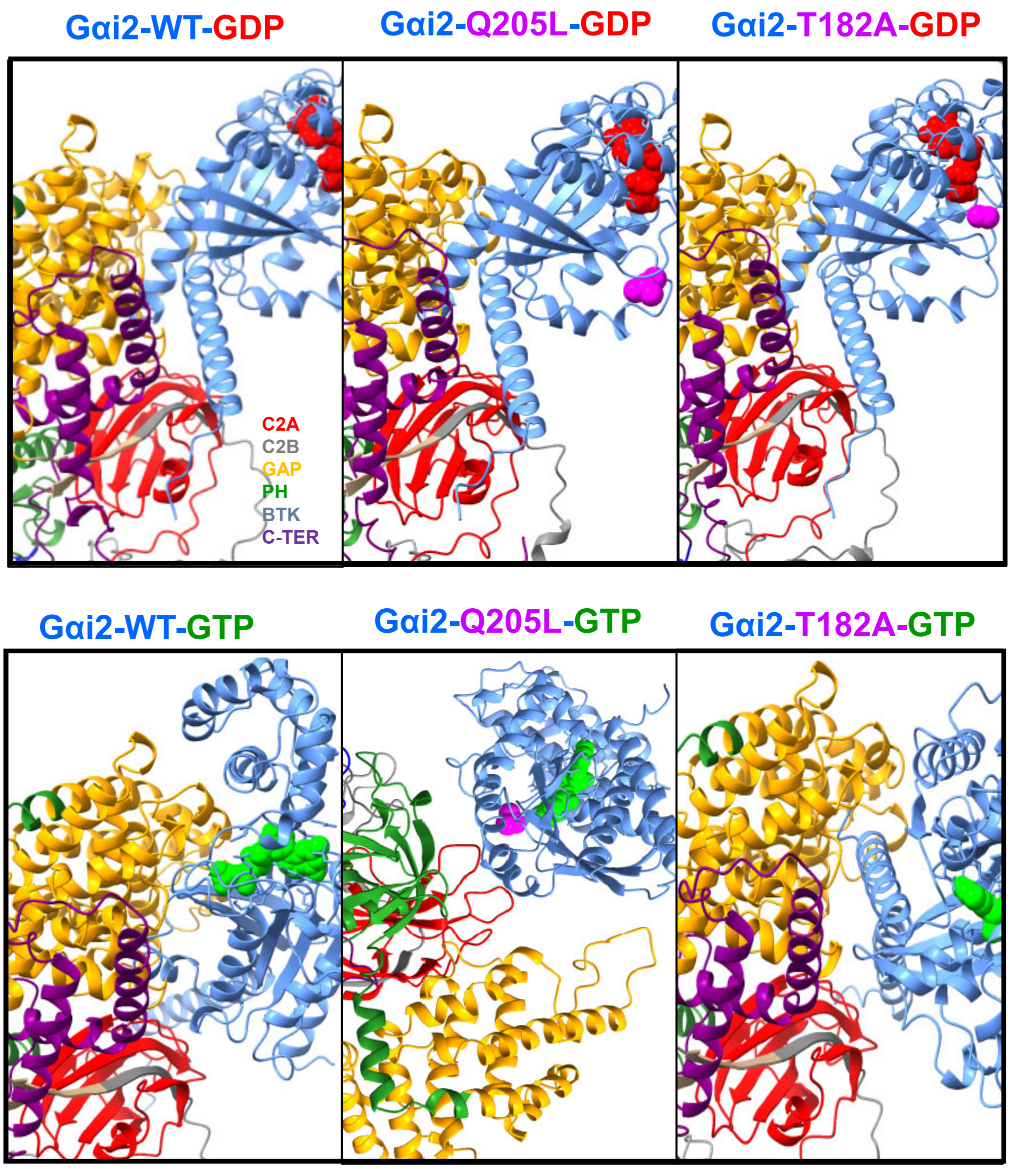


**B**

| | WT -GDP | WT -GTP | Q205L -GDP | Q205L -GTP | T182A -GDP | T182A -GTP |
|---|---|---|---|---|---|---|
| # of intermolecular contacts: | 121 | 131 | 99 | 94 | 131 | 134 |
| # of charged contacts: | 24.0 | 38.0 | 25.0 | 22.0 | 25.0 | 32.0 |
| # of charged-polar contacts: | 32.0 | 26.0 | 19.0 | 9.0 | 28.0 | 30.0 |
| # of charged-apolar contacts: | 33.0 | 28.0 | 18.0 | 39.0 | 42.0 | 30.0 |
| # of polar-polar contacts: | 4.0 | 8.0 | 5.0 | 1.0 | 5.0 | 7.0 |
| # of apolar-polar contacts: | 16.0 | 14.0 | 14.0 | 10.0 | 21.0 | 19.0 |
| # of apolar-apolar contacts: | 12.0 | 17.0 | 18.0 | 13.0 | 10.0 | 16.0 |
| % of apolar NIS residues: | 35.03 | 34.94 | 34.99 | 34.72 | 35.22 | 35.11 |
| % of charged NIS residues: | 33.763 | 4.023 | 34.10 | 33.79 | 33.60 | 33.37 |
| Predicted binding affinity (kcal/mol): | -13.2 | -12.7 | -11.1 | -12.8 | -15.1 | -13.7 |
| Predicated dissociation constant: | 2.2e-10 | 4.7e-10 | 7.5e-09 | 3.8e-10 | 8.8e-12 | 8.3e-11 |

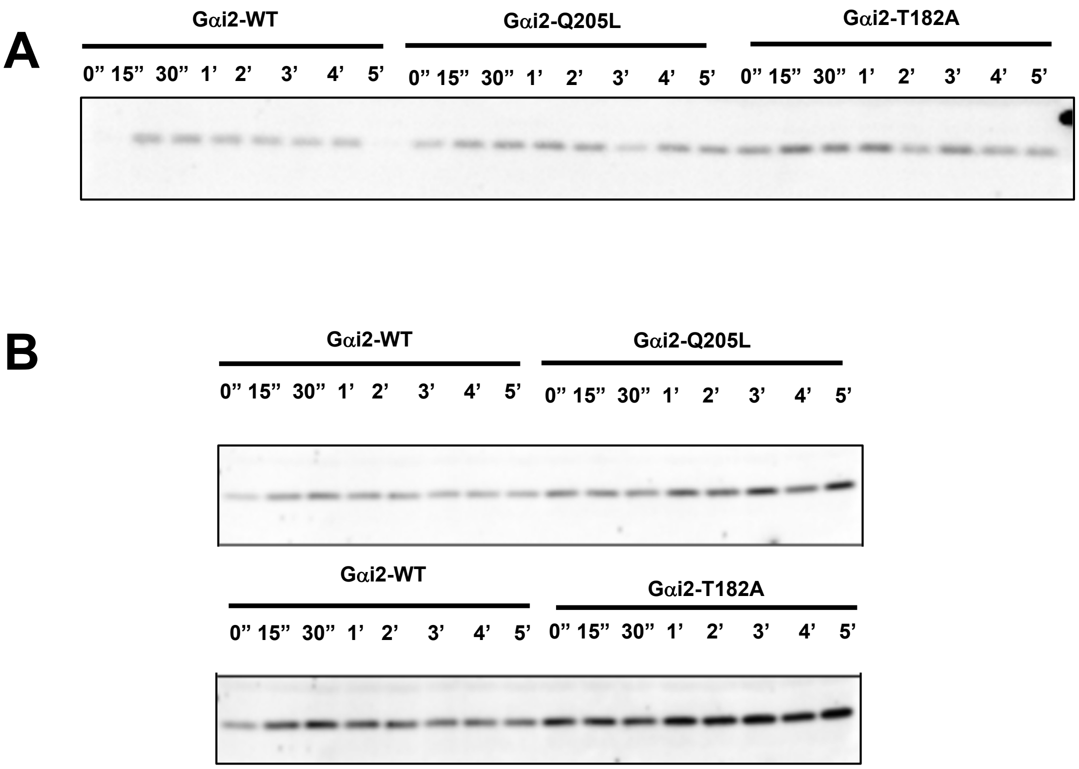
A
Gαi2-WT
Gαi2-Q205L
Gαi2-T182A
0” 15” 30” 1’ 2’ 3’ 4’ 5’ 0” 15” 30” 1’ 2’ 3’ 4’ 5’ 0” 15” 30” 1’ 2’ 3’ 4’ 5’
B
Gαi2-WT
Gαi2-Q205L
0” 15” 30” 1’ 2’ 3’ 4’ 5’ 0” 15” 30” 1’ 2’ 3’ 4’ 5’
Gαi2-WT
Gαi2-T182A
0” 15” 30” 1’ 2’ 3’ 4’ 5’ 0” 15” 30” 1’ 2’ 3’ 4’ 5’

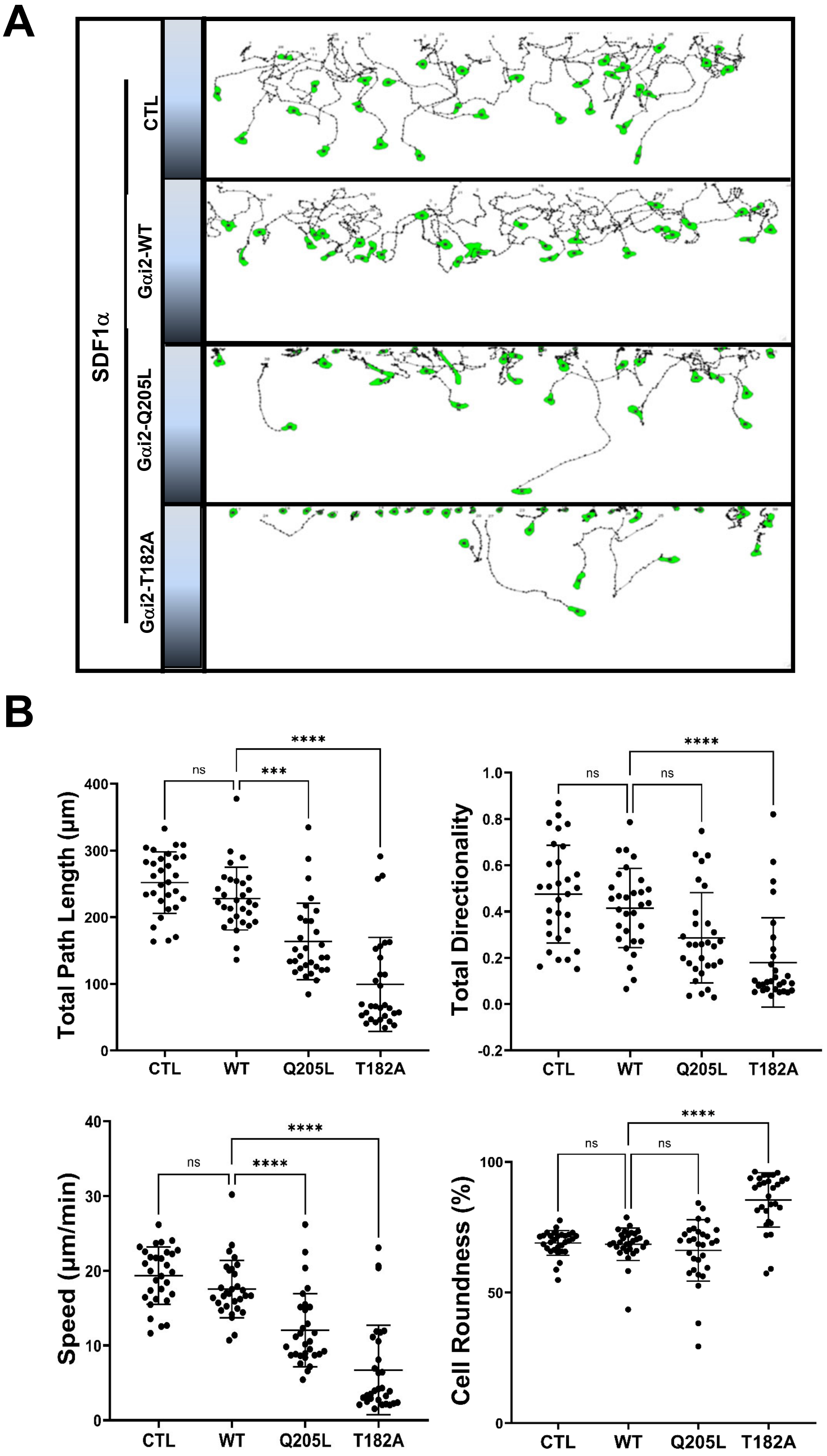
A
SDF1α
CTL
Gαi2-WT
Gαi2-Q205L
Gαi2-T182A
B
Total Path Length (μm)
Total Directionality
Speed (μm/min)
Cell Roundness (%)
ns
***
****
CTL
WT
Q205L
T182A

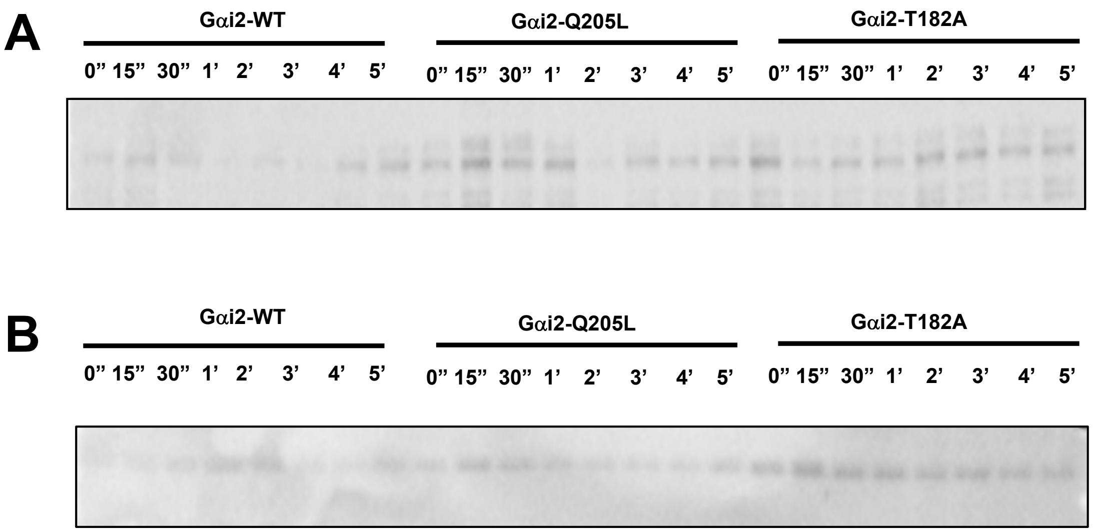
A
Gαi2-WT
Gαi2-Q205L
Gαi2-T182A
0" 15" 30" 1' 2' 3' 4' 5'
0" 15" 30" 1' 2' 3' 4' 5'
0" 15" 30" 1' 2' 3' 4' 5'
B
Gαi2-WT
Gαi2-Q205L
Gαi2-T182A
0" 15" 30" 1' 2' 3' 4' 5'
0" 15" 30" 1' 2' 3' 4' 5'
0" 15" 30" 1' 2' 3' 4' 5'
